# Innate–adaptive IFN cross-talk reprograms myeloid and T cells for antitumor immunity with a second-generation recombinant MVA

**DOI:** 10.1101/2022.09.25.509429

**Authors:** Shuaitong Liu, Yueqi Wang, Gregory Mazo, Shanza Baseer Tariq, Juan Angulo-Lozano, Ning Yang, Tuo Zhang, Yi Wang, Daniel Hirschhorn, Liangliang Ji, Adrian Tan, Jiahu Wang, Wei Yan, John Choi, Jenny Zhaoying Xiang, Ming O. Li, Taha Merghoub, Jedd D. Wolchok, Charles M. Rice, Liang Deng

## Abstract

Resistance to immune checkpoint blockade (ICB) remains a major obstacle in cancer immunotherapy. We rationally engineered a second-generation recombinant Modified Vaccinia virus Ankara (MQ833) by deleting viral immune-evasion genes (*E3L*, *E5R*, *WR199*) and incorporating *Flt3L*, *OX40L*, and matrix-anchored *IL12*. Intratumoral MQ833 elicited robust tumor regressions across multiple models, including ICB-resistant and MHC-I–deficient tumors. Single-cell RNA sequencing revealed extensive remodeling of the tumor microenvironment, characterized by neutrophil and monocyte recruitment and activation, M2 macrophage depletion, M1 polarization, and effector T-cell differentiation and proliferation. Conditional *Ifnar1* knockout mice demonstrated that MQ833 efficacy requires type I interferon signaling in neutrophils, macrophages/monocytes, and T cells. Moreover, *Nos2* deficiency impaired therapeutic efficacy, confirming iNOS⁺ myeloid cells as key effectors. Together, these findings show that MQ833 activates innate–adaptive IFN cross-talk to reprogram myeloid and T cells, defining a rationally designed viral immunotherapy capable of overcoming ICB resistance.

**GRAPHIC ABSTRACT:** 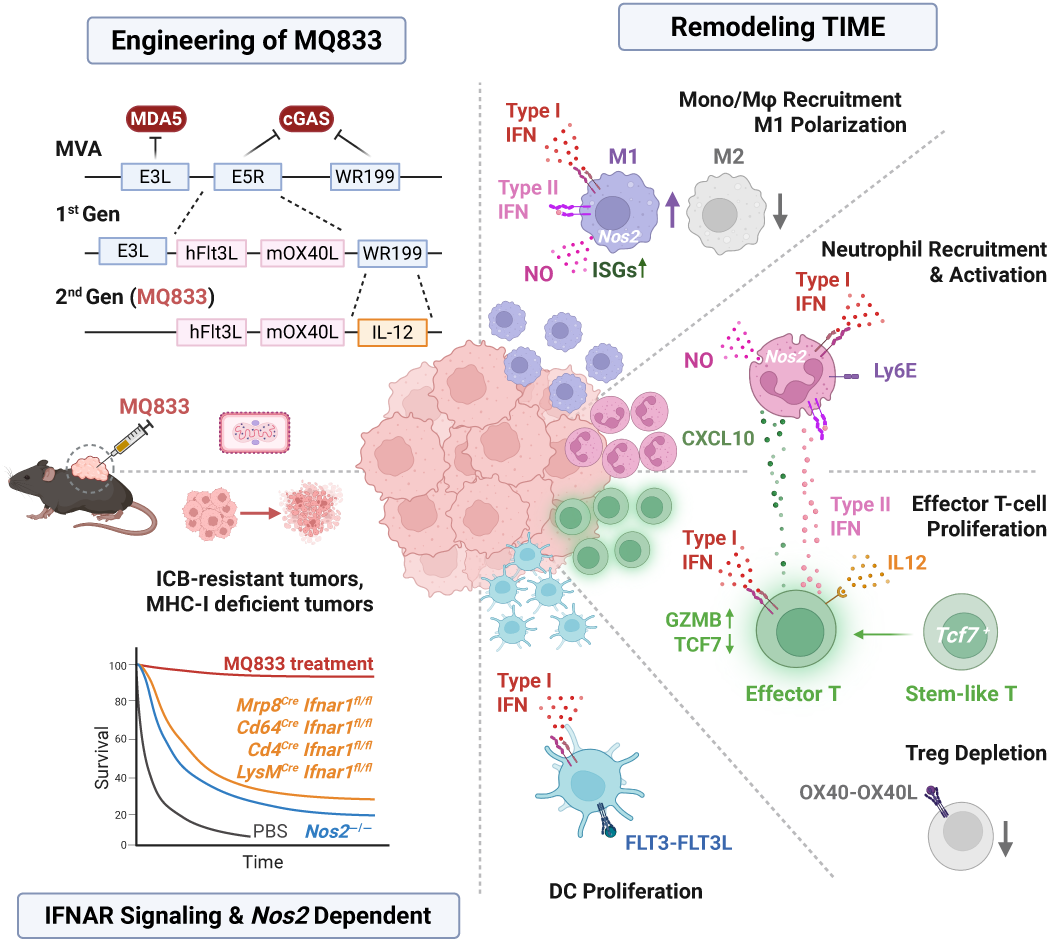

## INTRODUCTION

The tumor immune microenvironment (TIME) comprises diverse populations of myeloid and lymphoid cells whose antitumor functions are often subverted by the immunosuppressive milieu of growing tumors.^1–3^ A central challenge of cancer immunotherapy is to reprogram myeloid cells and reinvigorate T cells to sustain antitumor immunity.^4–7^ Among various approaches to overcome immune resistance, immunogenic viruses have emerged as potent tools to remodel the TIME and activate both innate and adaptive immunity.^8–11^ However, accumulating evidence suggests that viral replication and oncolysis are not essential for therapeutic efficacy; instead, activation of innate sensing and local inflammation are the dominant drivers of viral-induced antitumor immunity.^12–16^

Modified Vaccinia virus Ankara (MVA) is a large, non-replicative cytoplasmic DNA virus derived from the replication-competent vaccinia Ankara strain after more than 500 passages in chicken embryonic fibroblasts, resulting in extensive deletions in host range and immune evasion genes.^17,18^ MVA was recently approved as a vaccine against smallpox and mpox ^19^, and has been intensively studied as a vector for vaccines and cancer immunotherapies.^20,21^ Its favorable safety profile, large genomic capacity for transgene insertion, and ability to infect both tumor and immune cells make it an attractive backbone for rational vector design.

We previously showed that MVA infection of dendritic cells activates the cytosolic DNA-sensing pathway via cyclic GMP–AMP synthase (cGAS) and induces moderate type I interferon (IFN) production.^22^ Intratumoral (IT) injection of heat-inactivated MVA generates potent CD8⁺ T-cell–dependent antitumor immunity through cGAS–STING signaling.^14^ Furthermore, we demonstrated that heat-inactivated oncolytic vaccinia virus elicited stronger antitumor effects than its replicating counterpart,^15^ supporting the concept that immunogenic viral activation, rather than direct oncolysis, is sufficient to drive tumor regression. These findings established the basis for developing non-replicative, immunostimulatory MVA vectors for cancer immunotherapy.

Building on this foundation, we engineered a first-generation recombinant MVA (*rMVA*, MQ710) by deleting *E5R*, an inhibitor of cGAS, and inserting human *Flt3L* and murine *OX40L* to enhance dendritic-cell recruitment and regulatory T-cell depletion via OX40–OX40L and IFNAR signaling.^23^ Intratumoral delivery of MQ710 exhibited potent antitumor activity in murine models and is currently being evaluated in a Phase I clinical trial for advanced solid tumors (NCT05859074) with promising early results.

To further amplify immunogenicity and define the mechanistic basis of viral-induced antitumor immunity, we designed a second-generation rMVA, MQ833. MQ833 was engineered through deletion of three viral immune-suppressive genes (*E3L*, *E5R*, and *WR199*) and insertion of *Flt3L*, *OX40L*, and an extracellular matrix–anchored *IL12* transgene to enhance local cytokine activity while minimizing systemic toxicity. *E3L* encodes a double-stranded RNA–binding protein that suppresses MDA5-mediated sensing,^24,25^ whereas *WR199* inhibits cGAS activation;^26^ their removal was intended to potentiate both MDA5- and STING-dependent type I IFN signaling.

Here, we evaluated the therapeutic efficacy and mechanistic underpinnings of MQ833 across multiple murine tumor models, including melanoma, colon carcinoma, and bladder cancer. We used single-cell RNA sequencing to dissect how MQ833 reprograms the TIME, with particular focus on nucleic acid–sensing pathways and type I IFN–driven myeloid and T-cell activation. These studies define a mechanistically optimized recombinant platform that integrates rational viral engineering with localized cytokine delivery to orchestrate coordinated innate and adaptive immune activation against solid tumors

## RESULTS

### Rational engineering of MQ833 amplifies innate sensing and T cell activation

Our previous work identified the vaccinia E5R gene as a potent inhibitor of the DNA sensor cGAS. Deletion of E5R from MVA (MVAΔE5R) induced high levels of IFN-β in bone marrow-derived dendritic cells (BMDCs) in a cGAS-dependent manner.^26^ Based on this backbone, we generated our first-generation recombinant virus, rMVA (MQ710; MVAΔE5R-hFlt3L-mOX40L), which expresses membrane-bound human Flt3L and murine OX40L. Intratumoral (IT) delivery of MQ710 elicited strong antitumor immunity across multiple tumor models, dependent on CD8⁺ T cells, cGAS/STING, and IFNAR signaling, and also depleted intratumoral OX40^hi^ Tregs through OX40L–OX40 and IFNAR-mediated mechanisms.^23^

To further overcome viral immune evasion, we deleted additional inhibitory genes—E3L, which blocks cytosolic dsRNA sensing,^25^ and WR199 (MVA186), recently identified as a cGAS inhibitor.^27^ Deleting *E3L* together with *E5R* markedly increased *Ifnb* induction across multiple murine tumor cell lines, whereas single deletions yielded only modest effects (Figure S1A). Mechanistically, MVAΔE3LΔE5R triggered strong IRF3 phosphorylation and IFN-β production in WT but not in *Mda5⁻^/^⁻* or *Mda5⁻^/^⁻Sting⁻^/^⁻* tumor cells (Figure S1B and Figure S1C), confirming predominant dependence on MDA5 pathways.

In vivo, deletion of *E3L* enhanced tumor-specific T-cell priming. In a bilateral B16-F10 model, IT rMVAΔE3L induced a higher frequency of IFN-γ–secreting splenic CD8⁺ T cells compared with rMVA (Figure S2A-C). Further deletion of *WR199* produced MQ832 (MVAΔE5RΔE3LΔWR199-hFlt3L-mOX40L), which modestly increased CD8⁺ T-cell activation and further decreased OX40^hi^ Tregs compared with rMVAΔE3L (Figure S2D and Figure S2E).

To potentiate T/NK-cell activation while minimizing systemic cytokine toxicity, we inserted a bicistronic IL-12 cassette (Il12a–P2A–Il12b) under a modified H5 early/later promoter into the *WR199* locus and tagged the IL-12p35 subunit with an extracellular matrix-binding sequence to localize cytokine activity to the tumor microenvironment, generating MQ833 (MVAΔE5RΔE3LΔWR199-hFlt3L-mOX40L-mIL12) (Figure 1A and Figure 1B). ^28,29^ MQ833 infection induced robust transgene expression of hFlt3L and mOX40L, along with high levels of IL-12p40 secretion (Figure 1C and Figure 1D). MQ833 markedly upregulated *Ifnb* transcripts and IFN-β secretion in B16-F10 cells compared with MVA or MVAΔE5R (Figure 1E and Figure 1F). Western blot analysis showed enhanced STING and IRF3 phosphorylation beginning at 4 hours post infection (hpi), confirming strong activation of the cGAS/STING pathway (Figure 1G). Consistently, MQ833 induced high levels of STING-dependent IFN-β production and p-IRF3 in BMDCs from WT but not *Sting^Gt/Gt^* mice (Figure 1H and Figure 1I).

**Figure 1.**
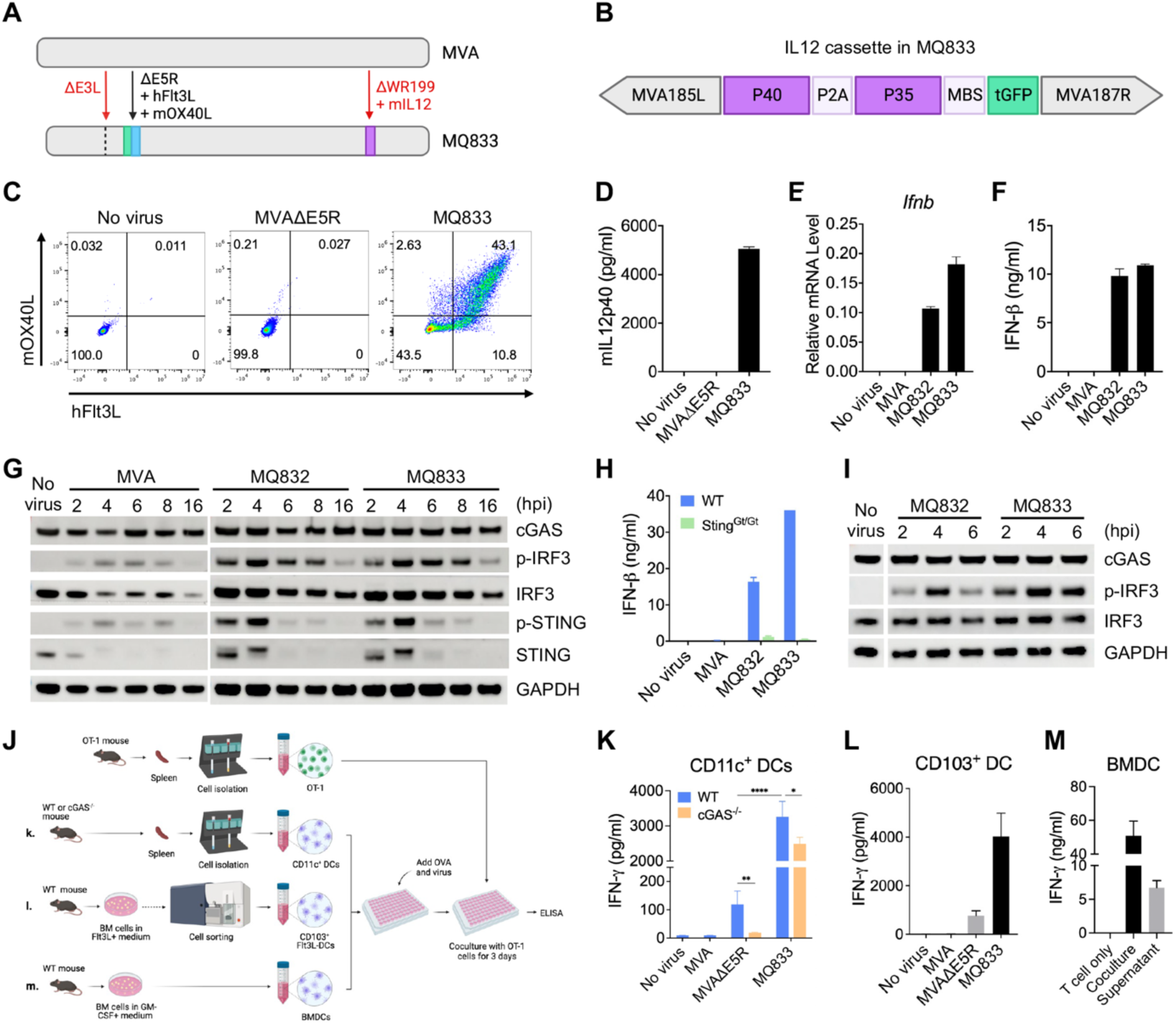
MQ833 infection of tumor and immune cells activates cGAS/STING mediated dsDNA sensing pathway and promotes IFN production. (A) Schematic of MQ833 design showing sequential deletions of viral immune-evasion genes (*E5R*, *E3L*, *WR199*) and insertion of transgenes encoding hFlt3L, mOX40L, and extracellular matrix-anchored mIL-12. (B) Diagram of the bicistronic IL-12 cassette (Il12a–P2A–Il12b) integrated into the *WR199* locus. (C and D) B16-F10 tumor cells infected with MVAΔE5R or MQ833 (MOI = 10) for 16 h. (C) Flow cytometry showing surface expression of hFlt3L and mOX40L. (D) ELISA quantifying secreted mIL-12p40. (E and F) B16-F10 tumor cells infected with indicated viruses (MOI = 10) for 16 h. qRT-PCR and ELISA showing increased *Ifnb* transcripts (E) and IFN-β secretion (F) in MQ833-infected cells versus controls. (G) Western blot analysis of cGAS/STING pathway activation in B16-F10 cells infected with indicated viruses (MOI = 10) and collected at 2–16 h post-infection (hpi). (H and I) BMDCs from WT or *Sting^Gt/Gt^* mice infected (MOI = 10). (H) ELISA quantification of IFN-β secretion. (I) Western blot for cGAS, p-IRF3, and IRF3. (J) Experimental schematic of DC–T-cell coculture used to evaluate antigen cross-presentation. (K–M) ELISA quantification of IFN-γ in supernatants from OT-I T-cell cocultures. (K) Splenic CD11c⁺ DCs from WT or *cGAS⁻^/^⁻* mice pulsed with OVA and infected with indicated viruses. (L) Sorted CD103⁺ DCs from Flt3L-expanded cultures pulsed with OVA and infected as in (K). (M) Comparison of OT-I activation using direct coculture versus conditioned medium from MQ833-infected BMDCs, showing contact-dependent T-cell stimulation. Data are representative of ≥ 2 independent experiments.

Functionally, MQ833 enhanced antigen cross-presentation and T-cell activation. Splenic CD11c⁺ DCs from WT or *cGAS⁻/⁻* mice infected with MQ833 and pulsed with OVA protein stimulated stronger OT-I T-cell IFN-γ responses than MVAΔE5R, with partial dependence on cGAS (Figure 1J and Figure 1K). Similar results were obtained with CD103⁺ DCs isolated from Flt3L-expanded BMDCs (Figure 1L). Conditioned medium from infected BMDCs induced minimal IFN-γ secretion compared with direct DC–T-cell coculture, indicating a contact-dependent mechanism (Figure 1M).

Finally, we evaluated a humanized version of MQ833 (hMQ833; MVAΔE5RΔE3LΔWR199-hFlt3L-hOX40L-hIL12) in human monocyte-derived dendritic cells (moDCs). Infection with hMQ833 led to strong hFlt3L/hOX40L surface expression and secretion of IL-12p70, accompanied by increased IFNB and CXCL10 transcription (Figure S3A-C). In co-culture assays, hMQ833-primed moDCs stimulated both allogeneic and autologous CD4⁺ and CD8⁺ T cells to secrete IFN-γ, with the most pronounced activation observed in allogeneic CD4⁺ T cells (Figure S3D and S3E).

Together, these results demonstrate that sequential deletion of viral immune evasion genes (*E5R*, *E3L*, *WR199*) synergistically enhances type I IFN induction through MDA5/STING signaling, while incorporation of extracellular matrix-bound IL-12 amplifies antigen cross-presentation and T-cell activation. MQ833 thus represents a second-generation recombinant MVA platform that coordinates innate–adaptive IFN cross-talk to overcome tumor immunosuppression.

### MQ833 induces superior antitumor activity through type I IFN and nucleic acid–sensing pathways

Having established that MQ833 infection of tumor and immune cells activates the cGAS/STING pathway and drives type I IFN production (Figure 1; Figures S1–S3), we next examined whether this enhanced innate sensing translates into superior antitumor efficacy in vivo. MQ833 differs from its predecessor MQ832 by incorporating an extracellular-matrix–anchored IL-12 cassette in addition to deletions of three viral immune-evasion genes (*E5R*, *E3L*, and *WR199*). This design was intended to further amplify local T-and NK-cell activation while minimizing systemic cytokine toxicity.

In the B16-F10 melanoma model, intratumoral (IT) administration of MQ833 resulted in complete tumor regression in all treated mice, whereas MQ832 produced partial responses with 40 % long-term survival (Figure 2A–2D). In the MC38 colon carcinoma model, MQ833 also outperformed MQ832, curing 100% of mice compared with 30% for MQ832 (Figures S4A– S4B).

**Figure 2.**
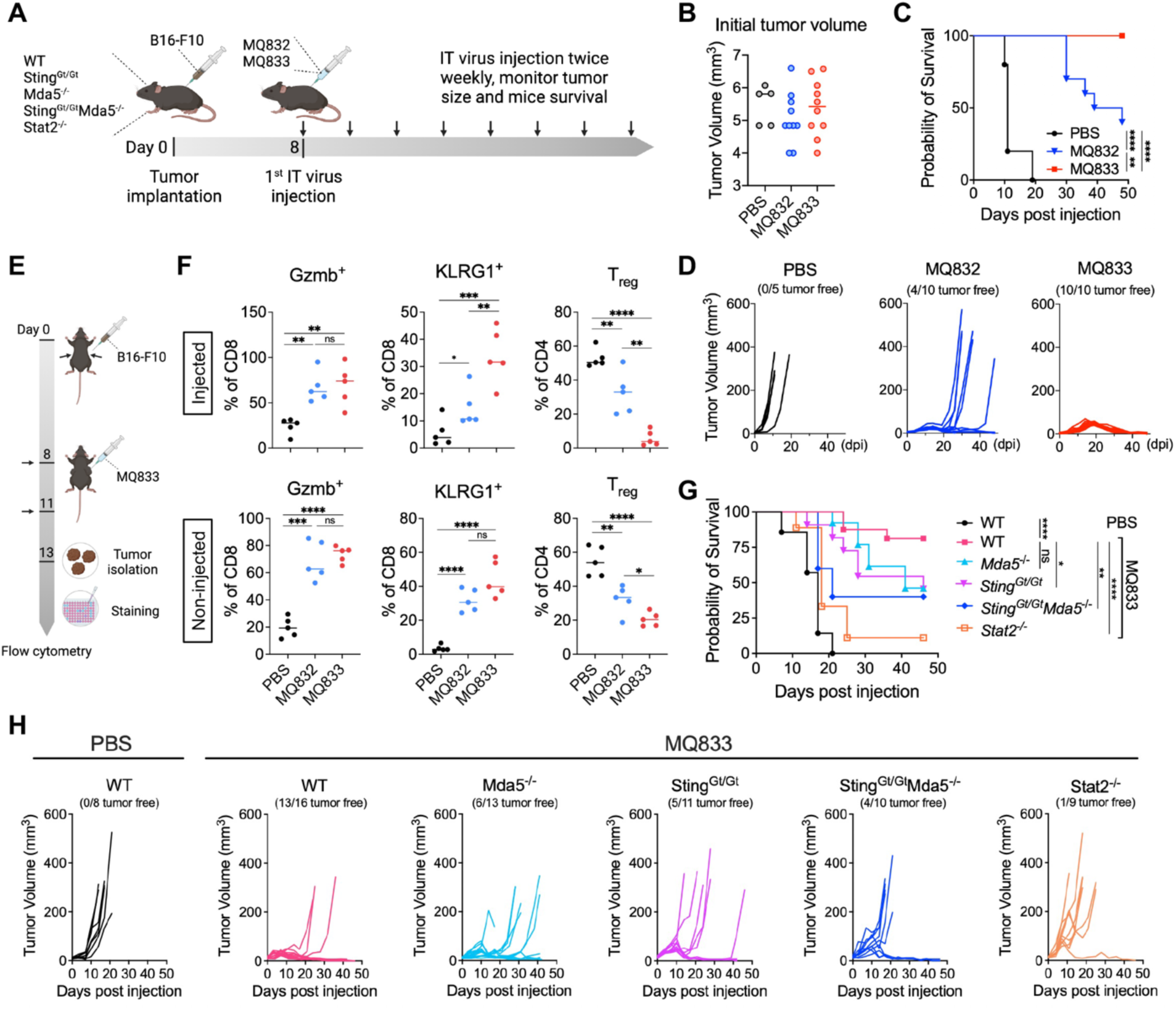
Intratumoral (IT) delivery of MQ833 generates an 80-100% cure in B16-F10 melanoma model in a STING-MDA5-STAT2-dependent manner. (A) Schematic diagram of survival experiment. (B) Kaplan-Meier survival curve of B16-F10 tumor bearing mice treated with MQ833. PBS was used as a control. (n=5-10) Survival data were analyzed by log-rank (Mantel-Cox) test. (*\*\*P<0.01, ****P < 0.0001*). (C) Initial tumor volume for each group. (D) Tumor volumes of mice injected with MQ832, MQ833 or PBS over days post treatment. (E) Schematic diagram of flow cytometric analysis of tumor infiltrating lymphocytes. (F) Percentage of specific cell population in the injected and non-injected tumors in different treatment groups. (G) Kaplan-Meier survival curve of B16-F10 tumor bearing mice treated with MQ833. PBS was used as a control. Survival data were analyzed by log-rank (Mantel-Cox) test. (n=5; *\*P<0.05, **P<0.01, ****P < 0.0001, t test*). (H) Tumor volume curves in different groups over days post treatment.

To assess systemic or abscopal effects, we next used a bilateral MC38 model with higher tumor burden, IT MQ833 delayed the growth of both injected and contralateral tumors and significantly prolonged survival relative to MQ832 (Figures S4C–S4E). Similar results were obtained in a bilateral B16-F10 model. We observed that IT MQ833 eradicated most injected tumors, markedly suppressed growth of contralateral non-injected lesions, and achieved cures in 40% of mice, whereas MQ832 produced no long-term survivors (Figures S5A–S5C). Combination therapy of IT MQ833 with systemic immune-checkpoint blockade (ICB) using anti-PD-1, anti-PD-L1, or anti-CTLA-4 antibodies further improved outcomes, curing 60 % of mice (Figures S5A–S5C). These findings demonstrate that MQ833 elicits potent local and systemic antitumor immunity that can be synergistically enhanced by ICB.

Flow-cytometric analyses of injected and non-injected tumors revealed that both MQ833 and MQ832 increased Granzyme B⁺ and KLRG1⁺ effector CD8⁺ T cells while reducing Foxp3⁺ regulatory T cells, but MQ833 induced a more pronounced and sustained Treg depletion across both tumor sites (Figure 2E–2F; Figures S5E–S5F).

Given its design to activate both dsRNA-and DNA-sensing pathways, we next evaluated MQ833 activity in mice deficient for *Mda5*, *Sting*, or both. IT MQ833 cured 81% of WT mice but only 45%, 46%, and 40% cure in *Sting*^Gt/Gt^, *Mda5*^−/−^, and *Sting*^Gt/Gt^*Mda5*^−/−^ mice, respectively (Figure 2G–2H), indicating that both MDA5- and STING-mediated nucleic-acid sensing contribute to its therapeutic efficacy. Furthermore, *Stat2⁻^/^⁻*mice responded poorly, with only 11% cures, confirming that type I IFN signaling through STAT2 is essential for MQ833-induced antitumor immunity (Figure 2G–2H).

Together, these results demonstrate that MQ833 engages dual nucleic-acid–sensing and type I IFN pathways to orchestrate coordinated innate and adaptive immune activation, resulting in complete and durable tumor regression in multiple preclinical models.

### MQ833 remodels myeloid and T-cell compartments in the tumor microenvironment

We next used single-cell RNA sequencing (scRNA-seq) to define how MQ833 alters the tumor immune microenvironment (TIME) and to assess the contributions of nucleic acid–sensing and type I IFN pathways. CD45⁺ immune cells were sorted from B16-F10 tumors in WT mice, *Sting*^Gt/Gt^*Mda5*^−/−^ (DKO), and Stat2^−/−^ mice two days after IT MQ833, followed by scRNA-seq analysis (Figure 3A). Unsupervised clustering identified 19 immune cell clusters, including T/NK cells, monocytes/macrophages, neutrophils, dendritic cells, and B cells, along with minor undefined and melanoma cell populations (Figure 3B and 3C). Cluster annotations were validated by expression of canonical lineage markers (Figure 3C; Figure S6A).

**Figure 3.**
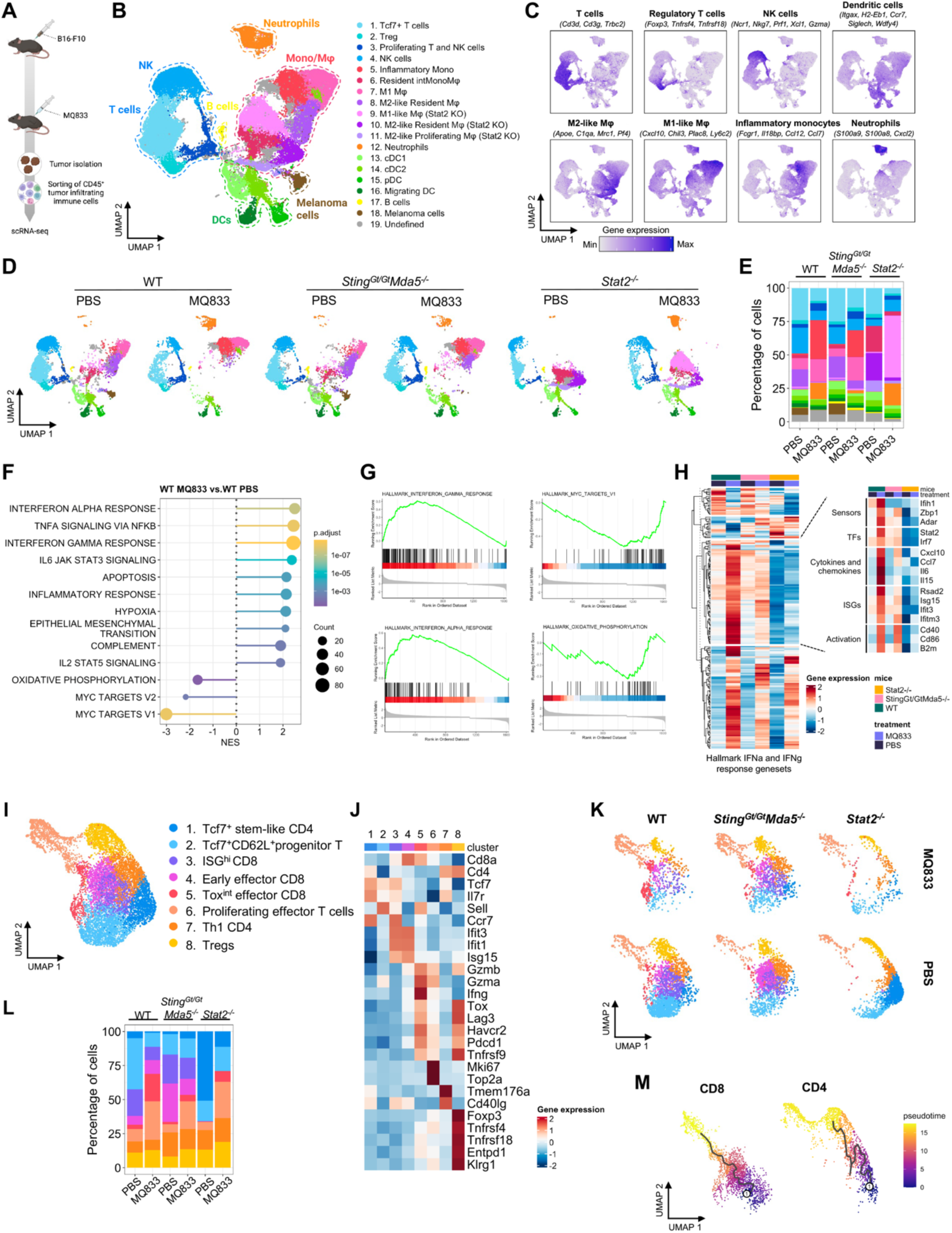
IT MQ833 treatment reprograms the tumor-infiltrated immune cells. (A) Schematic diagram of the experimental schedule. (B) UMAP display of sorted CD45^+^ cells from 6 samples combined following 10X Genomics scRNA-seq workflow (n = 47020 cells). (C) Expression plots of 8 main cell types using their top marker genes. (D) UMAP plots of cell clusters separated by sample. (E) Stacked bar plot showing the percentage of each cluster across different samples. (F) Dot plot for the top enriched and suppressed gene sets from the Molecular Signature Database (MSigDB) Hallmark immunologic signature gene sets in WT MQ833 versus WT PBS. (G) Gene set enrichment analysis (GSEA) for individual pathways from the representative enriched and suppressed gene sets in WT MQ833 sample compared with WT PBS. (H) Heatmaps of the average expression of genes in the hallmark IFNα and IFNγ response gene sets. Selected genes of interest are labeled in the right panel. (I) UMAP display of subclustered *Cd3e^+^* cells from 6 samples combined following 10X Genomics scRNA-seq workflow (n = 9,540 cells). (J) Cluster-defining marker expression was visualized in a heatmap. T cell subclusters includes Tcf7^+^ stem-like CD4 T cells (*Tcf7, Il7r, Cd4*), Tcf7^+^CD62L^+^ progenitor T cells (*Tcf7, Il7r, Sell*), ISG^hi^ CD8 T cells (*Tcf7, Ccr7, Ifit1, Ifit3, Isg15*), early effector CD8⁺ T cells (*Isg15, Ifit3, Gzmb, Ifng*), Tox^int^ effector CD8⁺ T cells (*Gzmb^hi^, Gzma^hi^, Ifng^hi^*), proliferating effector CD8⁺ T cells (*Mki67* and *Top2a)*. Th1 CD4⁺ T-cell (*Cd4, Il7r, Tmem176a*, *Cd40lg)*, and regulatory T cells (Tregs), marked by CD4, *Foxp3, Tnfrsf4* (OX40), *Tnfrsf18* (GITR), *Entpd1, Klrg1* expression. (K) UMAP plots of cell clusters separated by sample. (L) Stacked bar plot showing the percentage of each cluster across different samples. (M) Cell trajectory analysis of CD8 and CD4 T cells.

In WT mice, MQ833 induced rapid recruitment of inflammatory monocytes and neutrophils, which expanded from 1% to 29% and 0.3% to 12% of CD45⁺ cells, respectively (Figure 3D and 3E). This was accompanied by depletion of M2-like macrophages and increased M1-like macrophages, as well as reduced intratumoral cDC1 frequencies, consistent with migration to draining lymph nodes (Figure 3D and 3E). In DKO mice, neutrophil and monocyte recruitment and M2 depletion were attenuated, and cDC1 activation was impaired (Figure 3D and 3E). In *Stat2*⁻^/^⁻ mice, MQ833 triggered expansion of M1-like macrophage and neutrophil clusters that were transcriptionally distinct from their WT counterparts and lacked canonical IFN-stimulated gene (ISG) expression (Figure 3D–E). Single-cell viral transcript mapping revealed that neutrophils were the most heavily infected immune population (Figure S6B–S6C). Gene set enrichment analysis revealed robust induction of IFN-α, IFN-γ, and TNF signaling programs in MQ833-treated WT tumors, whereas oxidative phosphorylation and Myc targets were suppressed (Figure 3F and 3G). Heatmaps of IFN-responsive genes showed broad upregulation of ISGs, cytokines, and chemokines in WT mice, reduced responses in DKO mice, and complete loss in Stat2^−/−^ mice (Figure 3H).

Subclustering of CD3⁺ cells identified eight T-cell states: *Tcf7*^+^ stem-like CD4, *Tcf7*^+^*CD62L*^+^ progenitor T cell, ISG^+^ CD8, early effector CD8, *Tox^int^* effector CD8, proliferating effector T cells, Th1 CD4, and Tregs (Figure 3I and 3J). In WT mice, MQ833 decreased *Tcf7*⁺ stem-like subsets and expanded early effector, effector, and proliferating CD8⁺ T cells (Figure 3K and 3L). In DKO mice, differentiation into effector CD8⁺ subsets was attenuated, although proliferating CD8⁺ T cells still expanded. In mock-treated *Stat2*^−/−^ mice, *Tcf7*⁺ stem-like populations predominated, while effector subsets were absent. IT MQ833 in *Stat2*^−/−^ mice reduced stem-like populations and expanded proliferating CD8⁺ T cells, but failed to generate functional effectors, indicating that IL-12 alone cannot compensate for the absence of type I IFN signaling (Figure 3K and 3L). Pseudotime analysis revealed a continuous trajectory from stem-like to effector CD4⁺ and CD8⁺ T cells in WT tumors (Figure 3M).

Together, these results demonstrate that MQ833 reprograms the myeloid compartment by recruiting inflammatory monocytes and neutrophils, depleting M2-like macrophages, and promoting M1 polarization. Concurrently, MQ833 drives T-cell differentiation toward effector states in a manner dependent on MDA5/STING-mediated nucleic acid sensing and STAT2-dependent type I IFN signaling.

### Type I IFN–STAT2 signaling governs the transcriptional polarization, but not recruitment, of MQ833-induced tumor-infiltrating neutrophils

We next performed sub-clustering of tumor-infiltrating neutrophils to define their heterogeneity and to assess how type I IFN signaling shapes their activation states. Neutrophils were scarce at baseline but were robustly recruited after MQ833 treatment. UMAP analysis identified eight transcriptionally distinct neutrophil clusters (Figure 4A–4C). Clusters 1, 3, and 5 were enriched in WT mice following treatment and were characterized by high expression of interferon-stimulated genes (ISGs; *Ly6e, Isg15*) or *Nos2* (Figure 4D). The *ISG*^high^ *Ly6e*^high^ neutrophils represent antitumoral, pro-inflammatory subsets, consistent with prior findings that *Ly6e*^high^ neutrophils predict responsiveness to immunotherapy ^30^. Elevated *Nos2* expression further indicates tumoricidal potential through nitric-oxide-mediated cytotoxicity ^31^.

**Figure 4.**
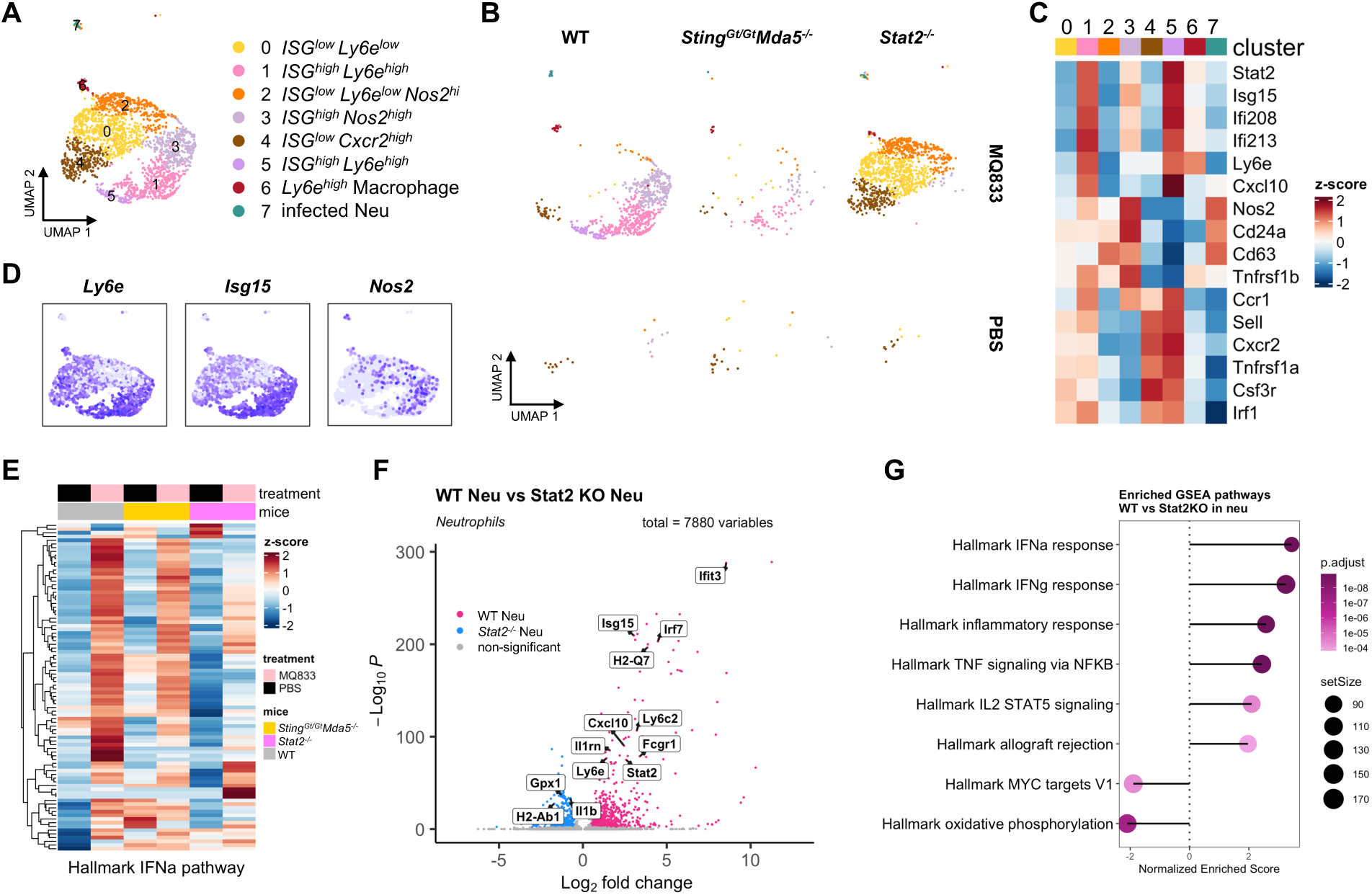
MQ833-induced type I IFN reprograms the tumor-infiltrating neutrophils. (A) UMAP display of subclustered neutrophils (cluster12 in figure. 3B) from 6 samples combined (n = 2,205 cells). (B) UMAP plots of cell clusters separated by sample. (C) Heatmap showing the expression of selected genes across different clusters. (D) Expression plots of *Ly6e*, *Isg15*, and *Nos2*. (E) Heatmaps of the average expression of genes in the hallmark IFNα response gene set. (F) Volcano plots comparing differentially expressed genes in neutrophils in WT and *Stat2^−/−^* mice. (G) Dot plot for the top enriched and suppressed gene sets from the Molecular Signature Database (MSigDB) Hallmark immunologic signature gene sets in MQ833-treated WT mice vs *Stat2^−/−^* mice.

Although *Sting*^Gt/Gt^*Mda5*^−/−^ mice showed detectable neutrophil infiltration after MQ833 treatment, the number and activation profiles of these cells were markedly reduced compared with WT tumors. In WT mice, robust neutrophil recruitment was accompanied by strong induction of *ISG*^high^ *Ly6e*^high^ and *Nos2*^high^ subsets, whereas in DKO mice, these populations were diminished and displayed attenuated expression of interferon-stimulated genes (Figure 4E). In addition, genes involved in the NF-kB pathway were also reduced in tumor-infiltrating CD45^+^ immune cells from DKO mice compared with WT mice (Figure 7SC). Thus, MDA5 and STING jointly promote both the recruitment and functional polarization of MQ833-induced tumor-infiltrating neutrophils.

In contrast, *Stat2*⁻^/^⁻ mice displayed transcriptionally divergent neutrophil clusters with markedly diminished expression of IFN-α–response genes (Figure 4E). Differential expression analysis (Figure 4F) revealed broad downregulation of canonical ISGs (*Ifit3, Isg15, Ly6c2, Ly6e, Stat2*), chemokines mediating T-cell recruitment (*Cxcl10*), and immune-modulatory genes (*Il1rn, Fcgr1, H2-Q7*). Pathway enrichment analysis (Figure 4G) demonstrated that MQ833-treated WT neutrophils showed activation of IFN-α/γ, inflammatory, and TNF–NF-κB signaling programs, whereas these pathways were blunted in *Stat2*⁻^/^⁻ mice. Conversely, MYC-target and oxidative phosphorylation programs were suppressed in WT but remained relatively enriched in *Stat2*⁻^/^⁻ neutrophils.

Collectively, these results demonstrate that MQ833 reprograms tumor-infiltrating neutrophils toward antitumoral and cytotoxic phenotypes, and that type I IFN–STAT2 signaling is indispensable for sustaining their pro-inflammatory transcriptional state and effector function.

### Intratumoral MQ833 remodels the tumor immune landscape locally and systemically in a bilateral melanoma model

To delineate the local and systemic immune effects of MQ833 treatment, we employed a bilateral B16-F10 melanoma model, in which the larger tumor received two intratumoral (IT) injections of MQ833 while the contralateral lesion remained untreated. Two days after the second injection, CD45⁺ cells were isolated from injected, non-injected, and PBS-treated control tumors and analyzed by single-cell RNA sequencing (Figure 5A). Unsupervised clustering identified 14 major immune populations, including T and NK cells, multiple macrophage and monocyte subsets, dendritic cells, neutrophils, and B cells (Figure 5B–C).

**Figure 5.**
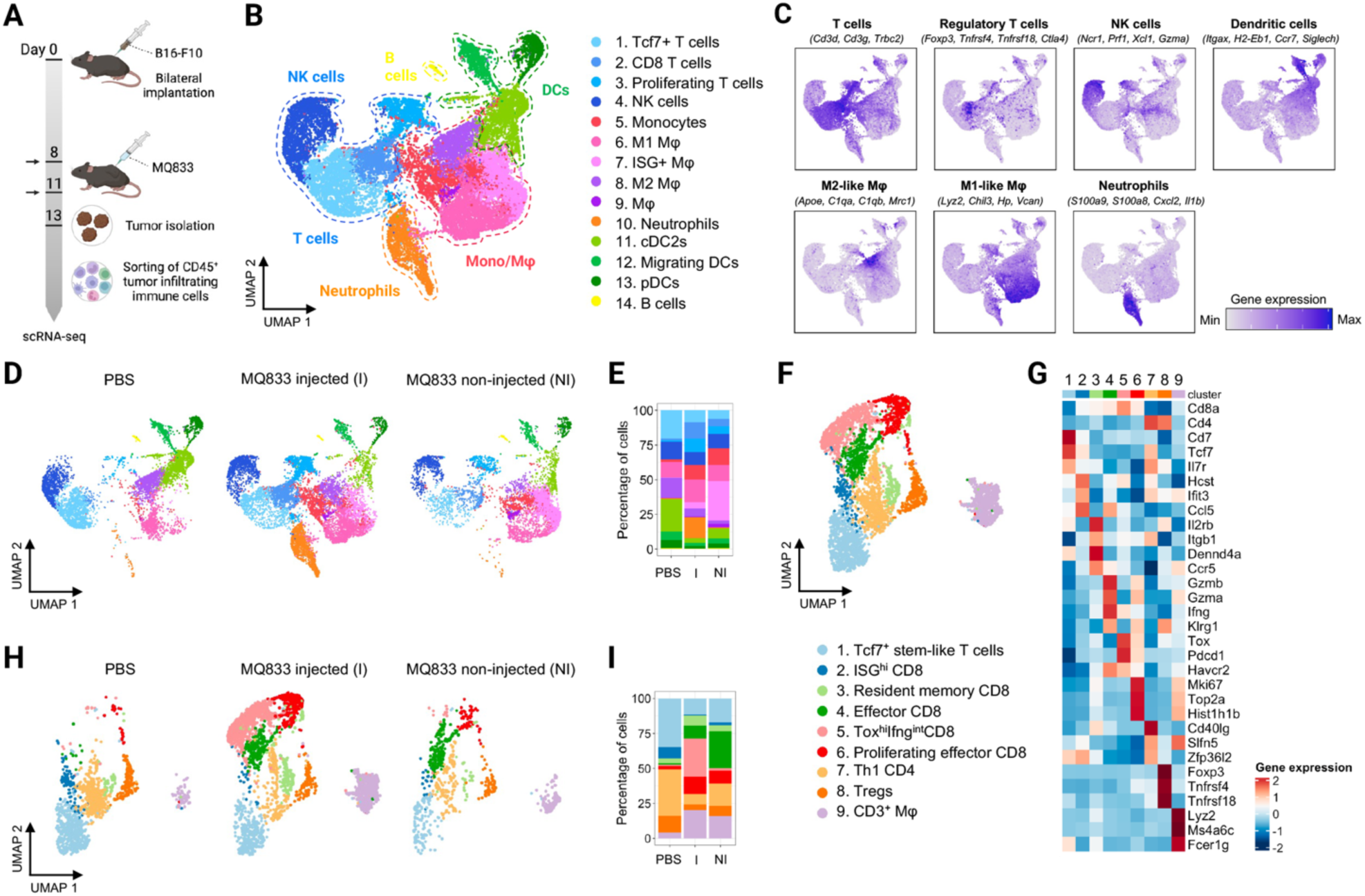
IT MQ833 reprograms TIME in both injected and non-injected tumors in a bilateral B16-F10 melanoma model. (A) Schematic diagram of the experimental schedule. (B) UMAP display of sorted CD45^+^ cells from 3 samples combined following 10X Genomics scRNA-seq workflow (n = 21,675 cells). (C) Expression plots of 7 main cell types using their top marker genes. (D) UMAP plots of cell clusters separated by sample. (E) Stacked bar plot showing the percentage of each cluster across different samples. (F) UMAP display of subclustered *Cd3e^+^* cells from 6 samples combined following 10X Genomics scRNA-seq workflow (n = 4,901 cells). (G) Heatmaps of the average expression of representative marker genes for each cluster. (H) UMAP plots of cell clusters separated by sample. (I) Stacked bar plot showing the percentage of each cluster across different samples.

IT MQ833 markedly reshaped the myeloid compartment (Figure 5D–E). Neutrophils, nearly absent in control tumors, became the dominant infiltrate in injected tumors but remained sparse in distant lesions, indicating localized recruitment. Inflammatory monocytes and M1-like macrophages expanded in both injected and non-injected tumors, while ISG⁺ macrophages were strongly enriched, particularly in the distant, non-injected tumors. In contrast, M2-like macrophages were depleted across both sites. These data indicate that MQ833 promotes an IFN- and inflammation-driven remodeling of the myeloid landscape, transforming suppressive myeloid populations into antitumor proinflammatory states.

Sub-clustering of CD3⁺ cells revealed nine transcriptionally distinct T-cell subsets (Figure 5F– G), including *Tcf7*^+^ stem-like T cells, ISG^+^CD8⁺, resident memory CD8^+^, effector CD8^+^, *Tox*^int^*Ifng*^int^ CD8, proliferating effector CD8, Th1 CD4⁺ T cells, Tregs, and a small CD3⁺ macrophage-like population. Following MQ833 treatment, effector, proliferating, and exhausted CD8⁺ T cells expanded markedly, particularly in the injected tumors, while ISG^+^CD8⁺, and stem-like subsets contracted. Effector CD8⁺ T cells also increased in the non-injected tumors, consistent with the abscopal effects observed at distant sites. In contrast, Th1 CD4⁺ T cells and Tregs were reduced, whereas CD3⁺ macrophage-like cells expressing *Lyz2*, *CD74*, *Ifitm3*, *Fcer1g*, Ms4a6c, *H2-Ab1*, *H2-Aa*, *Cxcl2*, increased at both tumor sites.

Collectively, these data show that IT MQ833 induces a dual-layered immune response. Locally, viral treatment triggers massive neutrophil recruitment, M1/ISG⁺ macrophage polarization, and proliferation of effector and exhausted CD8⁺ T cells, establishing a proinflammatory, cytotoxic milieu within injected tumors. Systemically, MQ833 promotes the expansion of ISG⁺ and M1 macrophages, accumulation of effector CD8⁺ T cells in distant, non-injected tumors—hallmarks of an abscopal, IFN-driven reprogramming of the tumor immune microenvironment.

### IFNAR signaling in both T cells and myeloid cells is required for the antitumor effects of MQ833

To define the cellular mediators of MQ833-induced tumor regression, we performed antibody-mediated depletion studies in the unilateral B16-F10 melanoma model (Figure 6A). Once tumors were established, mice received intratumoral MQ833 twice weekly together with systemic depletion of CD8⁺ or CD4⁺ T cells, NK1.1⁺ cells, Ly6G⁺ neutrophils, or CSF1-dependent macrophages. Flow-cytometric analysis of peripheral blood confirmed efficient and selective depletion for each target population (Figure S8A–C).

**Figure 6.**
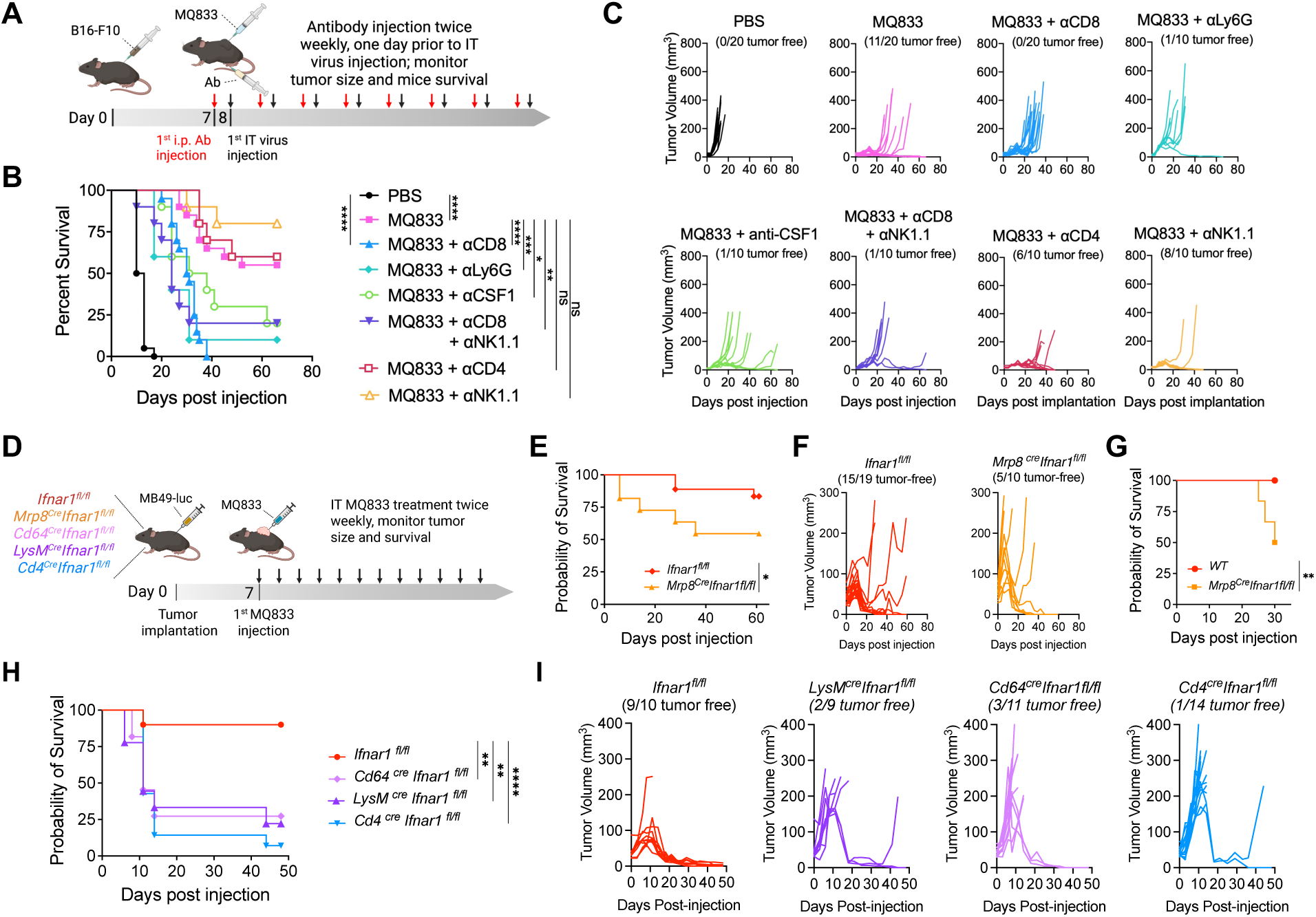
IFNAR signaling in both T cells and myeloid cells are important for the antitumor effects of MQ833. (A) Schematic diagram of the experimental schedule. (B) Kaplan-Meier survival curve of mice in each treatment group. (n=10) Survival data were analyzed by log-rank (Mantel-Cox) test. (n=10-20; *\*P < 0.05, **P < 0.01, ****P < 0.0001*). (C) Tumor volumes of mice separated by group over days post treatment. (D) Schematic diagram of the experimental schedule. (E) Kaplan-Meier survival curve of *Ifnar1^fl/fl^* and *Mrp8^Cre^Ifnar1^fl/fl^* mice. Survival data were analyzed by log-rank (Mantel-Cox) test. (n=10-19; *\*P < 0.05*). (F) Tumor volumes separated by group over days post treatment. (G) Kaplan-Meier survival curve of rechallenged mice in each group. Survival data were analyzed by log-rank (Mantel-Cox) test. (n=10-14; *\*\*P < 0.01*). (H) Kaplan-Meier survival curve of *Ifnar1^fl/fl^*, *Cd64^Cre^Ifnar1^fl/fl^*, *LysM^Cre^Ifnar1^fl/fl^*, *and Cd4^Cre^Ifnar1^fl/fl^*. Survival data were analyzed by log-rank (Mantel-Cox) test. (n=9-14; *\*P < 0.05, ****P < 0.0001*). (I) Tumor volumes of mice separated by group over days post treatment.

IT MQ833 alone achieved durable tumor clearance in more than half of treated animals, whereas CD8⁺ T-cell depletion completely abolished therapeutic benefit, confirming their indispensable role in viral immunotherapy (Figure 6B–C). Strikingly, depletion of neutrophils or macrophages also markedly reduced tumor control and survival, demonstrating that myeloid cells are essential contributors to MQ833-driven antitumor efficacy. In contrast, depletion of CD4⁺ T cells or NK cells had minimal impact on early tumor regression (Figure 6B–C). Together, these data establish that MQ833 engages a cooperative network of cytotoxic CD8⁺ T cells, neutrophils, and macrophages to mediate its potent therapeutic effects.

Because type I interferons orchestrate antiviral and antitumor inflammation, we next asked whether IFNAR signaling in distinct immune compartments mediates MQ833-driven tumor control. MB49-luc bladder tumors were implanted into conditional *Ifnar1^fl/fl^* mice lacking IFNAR selectively in *Mrp8⁺* neutrophils, *Cd64⁺* macrophages, *LysM⁺* myeloid cells, or *Cd4⁺* T cells (Figure 6D). IT MQ833 achieved 79% survival in *Ifnar1^fl/fl^* controls but only 50% in *Mrp8^Cre^Ifnar1^fl/fl^* mice, indicating that neutrophil-intrinsic IFNAR signaling contributes substantially to antitumor efficacy (Figure 6E–F). Notably, all *Ifnar1^fl/fl^* mice that achieved tumor clearance following MQ833 treatment resisted subsequent tumor rechallenge, whereas 60% of *Mrp8^Cre^Ifnar1^fl/fl^* mice failed to reject (Figure 6G), indicating that intact IFNAR signaling in neutrophils is required not only for primary tumor control but also for the establishment of durable antitumor memory. Loss of *Ifnar1* in *Cd64⁺* or *LysM⁺* myeloid cells similarly compromised tumor rejection, underscoring the importance of myeloid-intrinsic type I IFN signaling (Figure 6H-I). Deletion of *Ifnar1* in *Cd4⁺* T cells, which eliminates IFNAR1 expression in both CD4⁺ and CD8⁺ T-cell compartments, also partially reduced efficacy (Figure 6H–I), indicating that type I IFN signaling within T cells is critical for sustaining cytotoxic effector differentiation and long-term antitumor immunity.

Collectively, these findings demonstrate that MQ833-induced tumor regression requires coordinated IFNAR signaling across myeloid and T cells, which together orchestrate the inflammatory and effector programs necessary for durable tumor eradication.

### Nos2-dependent myeloid activation contributes to MQ833-induced antitumor immunity

To determine whether MQ833 engages nitric oxide–mediated effector mechanisms, we analyzed tumor-infiltrating immune cells two days after intratumoral injection. Flow cytometry revealed a marked increase in iNOS⁺ (Nos2⁺) neutrophils and M1-like macrophages, accompanied by a decrease in CD206⁺ M2-like macrophages (Figure 7A–C), indicating that MQ833 polarizes the myeloid compartment toward a proinflammatory phenotype.

**Figure 7.**
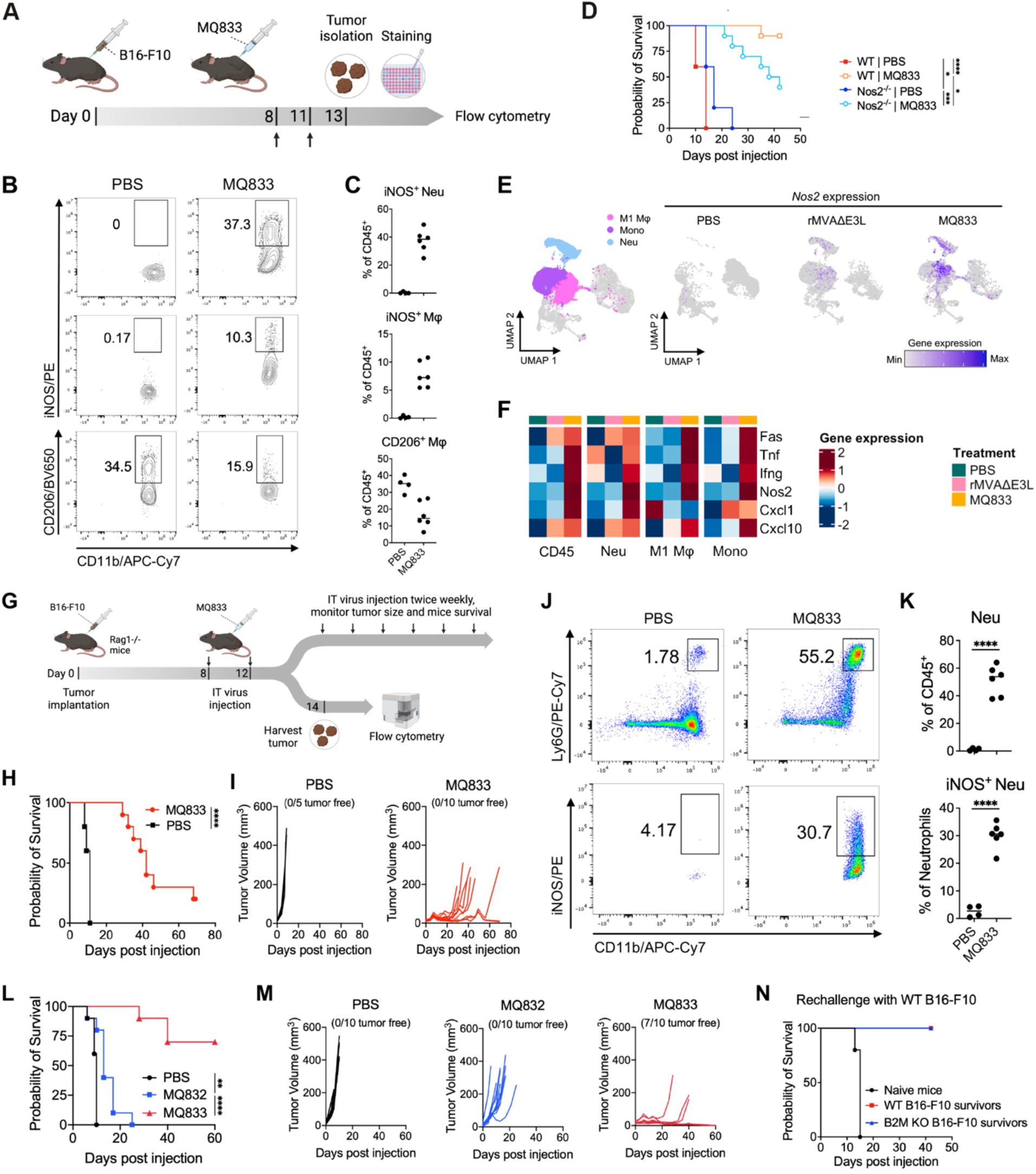
*Nos2*-dependent myeloid activation contributes to MQ833-induced antitumor immunity. (A) Schematic diagram of the flow cytometry analysis experiment. (B and C) Contour plots (B) and dot plots (C) showing percentage of iNOS^+^ cells among CD45^+^CD3^−^CD11b^+^Ly6g^+^ neutrophils (top row), CD45^+^CD3^−^CD68^+^CD11b^+^ macrophages (middle row), and percentage of CD206^+^ cells among macrophages (bottom row) in MQ833 or PBS treated tumors. (D) Kaplan-Meier survival curve of mice in each treatment group. Survival data were analyzed by log-rank (Mantel-Cox) test. (n=5-10; *\*P < 0.05, ***P < 0.001, ****P < 0.0001*) (E and F) scRNA-seq analysis of CD45^+^ cells isolated from rMVAΔE3L or MQ833 treated tumors. (E) UMAP display showing *Nos2* expression (right panel) in highlighted clusters (left panel) (F) Heatmaps showing average expression of selected antitumor marker genes in all the cells (first panel), neutrophil (second panel), M1 macrophage (third panel), and monocyte (fourth panel) clusters divided by treatment groups. (G) Schematic diagram of the survival and flow cytometry experimental design. (H) Kaplan-Meier survival curve of mice in each treatment group. PBS was used as a control. Survival data were analyzed by log-rank (Mantel-Cox) test. (n=5-10; *\*\*\*\*P < 0.0001*). (I) Tumor volumes of mice separated by group over days post treatment. (J and K) Dot plots (D) and bar graphs (E) showing percentage of CD45^+^CD3^−^CD11b^+^Ly6g^+^ neutrophils among total CD45^+^ cells (top row) and iNOS^+^ cells among neutrophils (bottom row) in MQ833 or PBS treated tumors. (L and M) 1×10^6^ of B2m^−/−^ B16-F10 cells were intradermally implanted into the right flank of WT C57BL/6J mice. (L) Kaplan-Meier survival curve of mice in each treatment group. PBS was used as a control. (n=10) Survival data were analyzed by log-rank (Mantel-Cox) test. (n=10; *\*\*P < 0.01, ****P < 0.0001*). (M) Tumor volumes of mice separated by group over days post treatment. (N) Mice rejected primary B16-F10 or B2m^−/−^ B16-F10 tumors were rechallenged with a lethal dose of B16-F10 tumor cells. Kaplan-Meier survival curve of the rechallenged mice.

We next evaluated whether Nos2 expression is required for the therapeutic activity of MQ833. In WT mice bearing B16-F10 tumors, IT MQ833 resulted in 90% survival rate in WT mice and only 40% survival rate in *Nos2*^−/−^ mice (Figure 7D), establishing a key role for nitric oxide in mediating tumor clearance.

Single-cell RNA-seq profiling of CD45⁺ tumor-infiltrating cells showed that both MQ833 and rMVAΔE3L expanded inflammatory monocytes and neutrophils while reducing M2 macrophages, but MQ833 uniquely upregulated *Nos2*, *Fas*, *Tnf*, *Cxcl1*, and *Cxcl10* in myeloid clusters (Figure 7E–F; Figure S9A–C). Subclustering of CD3⁺ cells further revealed that MQ833 promoted effector and effector-memory CD8⁺ T cells (C5, C7) while depleting Tcf7⁺ stem-like subsets (Figure S9D–G). These coordinated changes reflect IL-12–driven CD8⁺ T-cell proliferation, activation, and IFN-γ production, which in turn induce *Nos2*⁺ myeloid cells. CXCL10 produced by these activated myeloid cells further enhances recruitment of effector T cells into inflamed tumors.

Furthermore, MQ833 retained partial activity in *Rag1⁻^/^⁻* mice lacking T and B cells, prolonging survival despite the absence of adaptive immunity (Figure 7G–I). Tumors from these mice showed strong infiltration of iNOS⁺ neutrophils (Figure 7J–K), confirming that MQ833 can mobilize innate cytotoxic programs independently of lymphocytes.

Finally, MQ833 was highly effective against *B2m⁻^/^⁻* B16-F10 tumors that lack MHC I and are resistant to T-cell killing. IT MQ833 eradicated 70% of *B2m⁻^/^⁻* tumors, whereas control virus MQ832 produced no cures (Figure 7L–M). All cured mice rejected rechallenge with WT B16-F10 cells (Figure 7N), demonstrating durable systemic immunity.

Together, these results define a dual mechanism by which MQ833 orchestrates IL-12–induced, IFN-γ–dependent activation of Nos2⁺ myeloid cells, which in turn synergize with adaptive T-cell immunity to eliminate tumors, including that refractory to checkpoint blockade.

## DISCUSSION

Viral-based cancer immunotherapy can remodel the tumor immune microenvironment (TIME) to overcome both primary and acquired resistance to immune checkpoint blockade (ICB). Infection of tumor and immune cells by recombinant viruses activates innate immune sensing, induces immunogenic cell death, and promotes antigen-specific T-cell priming. Here, we describe the rational engineering of a recombinant Modified Vaccinia virus Ankara (rMVA), MQ833, through sequential deletion of three immune-evasion genes (*E3L*, *E5R*, and *WR199*) and insertion of three immunostimulatory transgenes (*Flt3L*, *OX40L*, and *IL-12*), thereby amplifying MDA5–STING–STAT2–dependent innate sensing and coordinating myeloid and T-cell activation to elicit potent antitumor immunity.

Antibody-depletion studies established that CD8⁺ T cells, neutrophils, and macrophages are indispensable for MQ833-mediated tumor regression. Using genetically engineered mice deficient in nucleic-acid sensing or type I IFN signaling, we showed that MQ833 requires MDA5/STING-driven cytosolic sensing and STAT2-dependent type I IFN signaling for durable tumor control. Single-cell RNA sequencing (scRNA-seq) revealed that intratumoral MQ833 reprograms the TIME, driving inflammatory myeloid polarization and effector T-cell differentiation. These results define MQ833 as a mechanistically engineered viral platform that rewires innate–adaptive immunity to overcome immune evasion.

MQ833 infection triggers robust type I IFN and pro-inflammatory cytokine production through MDA5/STING activation. Localized expression of extracellular-matrix-anchored IL-12 promotes differentiation of TCF7⁺ stem-like T cells into effector populations, augments IFN-γ production, and polarizes macrophages toward an M1 phenotype. Concurrently, MQ833 recruits neutrophils, where IFNAR/IFNGR signaling induces Nos2 expression and nitric oxide–mediated cytotoxicity, transforming an immunosuppressed milieu into an IFN-driven cytotoxic network.

MQ833 builds upon the first-generation rMVA (MVAΔE5R-hFlt3L-mOX40L) by deleting E3L, a dsRNA-binding protein that blocks MDA5 sensing, and WR199, a cGAS inhibitor. Combined deletion of E5R and E3L synergistically enhanced type I IFN induction, while addition of IL-12 yielded maximal immune activation. This demonstrates that iterative, mechanism-based vector design can amplify innate recognition without compromising viral stability.

Localized IL-12 expression increased effector CD8⁺ T cells, reduced intratumoral Tregs, and enhanced IFN-γ production in melanoma and colon-carcinoma models. scRNA-seq showed expansion of effector CD8⁺ T cells and upregulation of *Nos2*, *Fas*, *Tnf*, and *Cxcl10* in myeloid clusters, indicating enhanced cytotoxic and chemotactic capacity. Efficacy was reduced in *Nos2*⁻^/^⁻ mice, establishing nitric oxide as a key effector linking innate and adaptive immunity and underscoring that IL-12 and type I IFN signaling act in concert to sustain effector programs.

In addition to T-cell–mediated effects, MQ833 displayed CD8⁺ T-cell–independent activity through neutrophils and macrophages. Depletion of Ly6G⁺ neutrophils or CSF1-dependent macrophages abrogated efficacy, while MQ833 delayed tumor growth in *Rag1*⁻^/^⁻ mice and eradicated MHC I–deficient tumors, indicating that MQ833 mobilizes both innate and adaptive effector programs—distinguishing it from prior oncolytic viruses dependent on T-cell killing.

Neutrophils displayed a prominent IFN-stimulated-gene (ISG) signature after viral treatment. scRNA-seq identified *Ly6E^high^ ISG^high^* and *Nos2^high^* subsets consistent with antitumoral neutrophil states. These subsets were diminished in *Mda5*⁻^/^⁻ *Sting^Gt/Gt^* mice and absent in *Stat2*⁻^/^⁻ mice, demonstrating that type I IFN–STAT2 signaling governs neutrophil polarization. MQ833-infected neutrophils upregulated Cxcl10, linking them to IFN-driven T-cell recruitment and defining a neutrophil–T-cell cross-talk axis critical for viral immunotherapy.

Recent studies revealed that interferon-stimulated neutrophil states predict immunotherapy outcomes. IFN-licensed neutrophils cooperate with T cells to eliminate antigen-escape variants,^31^ while conserved *ISG*⁺*Nos2*⁺ or *Ly6E*⁺ subsets driven by tumor-intrinsic STING correlate with tumor control.^30,32^ The MQ833-induced Ly6E⁺ISG⁺ neutrophils likely represent this IFN-conditioned effector subset, bridging innate and adaptive immunity through coordinated IFNAR– STAT2 and IL-12–IFN-γ–NOS2 signaling. These findings position neutrophil-intrinsic IFN signaling as a key axis of antitumor immunity and expand the effector landscape of viral-based cancer immunotherapy. Consistent with these observations, our conditional *Ifnar1* deletion models confirm that IFNAR signaling in neutrophils and other myeloid lineages is indispensable for MQ833 efficacy.

Loss of IFNAR1 in *Mrp8*⁺ neutrophils, *Cd64*⁺ macrophages, *LysM*⁺ myeloid cells, or *Cd4*⁺ T cells each attenuated tumor regression, underscoring the cooperative requirement for IFNAR signaling across immune compartments. Neutrophil-specific *Ifnar1* deletion (*Mrp8^Cre^Ifnar1^fl/fl^*) impaired tumor control and memory formation, confirming that neutrophil-intrinsic IFNAR signaling contributes to both cytotoxicity and long-term protection. Similarly, macrophage- or T-cell-specific IFNAR loss reduced efficacy, demonstrating that MQ833-induced tumor clearance relies on coordinated IFNAR–STAT2 signaling across innate and adaptive lineages.

These results extend classical paradigms of type I IFN immunity. Earlier studies showed that Ifnar1⁻^/^⁻ mice fail to reject immunogenic tumors,^33^ and that IFNAR signaling in dendritic cells is required for CD8⁺ T-cell cross-priming and tumor rejection,^34,35^ while tumor cell–intrinsic IFNAR promotes antigen presentation and immunogenic cell death.^36^ Our findings advance this framework by revealing that, during viral immunotherapy, IFNAR signaling in macrophages, neutrophils, and T cells is equally essential for tumor regression and memory formation, highlighting a coordinated innate–adaptive IFN network as the central mechanism of viral-induced antitumor immunity.

The abscopal effects of MQ833 underscore its systemic reach. In bilateral tumor models, intratumoral MQ833 reshaped both injected and distant lesions, expanding M1/ISG⁺ macrophages and effector CD8⁺ T cells while reducing Tregs and M2 macrophages. These results demonstrate that local viral infection propagates systemic immune activation through type I IFN–dependent circuits that synchronize cytotoxic activity across tumor sites.

Mechanistically, MQ833 differs from lytic oncolytic viruses such as HSV-1–derived T-VEC ^37^ or adenovirus CG0070.^38^ Whereas T-VEC primarily achieves local tumor lysis and induces regional immune activation with modest systemic effects, MQ833 amplifies cytosolic nucleic acid sensing and IFN-driven myeloid–T-cell cross-talk through rational gene deletions and localized IL-12 expression. This strategy achieves durable systemic immunity with minimal toxicity, exemplifying a paradigm shift from direct oncolysis to immune orchestration.

From a translational perspective, MQ833 provides a versatile framework for rational viral design. Its ability to eradicate MHC I–deficient and ICB-resistant tumors highlight myeloid activation as a complementary mechanism of tumor control. Because MVA-based platforms are well-characterized, scalable, and clinically proven to be safe, the design principles established here can be rapidly applied to clinical-grade constructs.

In summary, MQ833 is a second-generation recombinant MVA that integrates rational viral engineering with localized cytokine delivery to achieve robust, multifaceted antitumor immunity. By combining deletion of viral immune-evasion genes (*E3L*, *E5R*, *WR199*) with extracellular-matrix-anchored IL-12 expression, MQ833 activates dual MDA5–STING sensing and amplifies STAT2-mediated type I IFN signaling within the tumor microenvironment. These coordinated pathways remodel the myeloid and T-cell compartments, driving recruitment and activation of *Nos2*⁺ neutrophils and M1-polarized macrophages, depletion of Tregs, and expansion of effector-memory CD8⁺ T cells.

The discovery that myeloid and T cell-intrinsic IFNAR signaling underlies tumor control and antitumor memory redefines the effector architecture of viral immunotherapy, positioning MQ833 as a mechanistically defined and clinically translatable platform for combination immunotherapy.

## METHODS

### Mice

Female C57BL/6J mice between 6 and 8 weeks of age were purchased from the Jackson Laboratory (Strain #000664) were used for the preparation of bone marrow-derived dendritic cells and for *in vivo* experiments. These mice were maintained in the animal facility at the Sloan Kettering Institute. All procedures were performed in strict accordance with the recommendations in the Guide for the Care and Use of Laboratory Animals of the National Institutes of Health. The protocol was approved by the Committee on the Ethics of Animal Experiments of Sloan Kettering Cancer Institute. *Sting^Gt/Gt^* mice were generated in the laboratory of Dr. Russell Vance (University of California, Berkeley).^39^ *cGas^−/−^*(Strain #026554),^40^ *Stat2^−/−^* (Strain #023309)^41^ and *Nos2*^−/−^ (Strain #002609)^42^ mice were purchased from Jackson Laboratory. *Mda5*^−/−^ mice were generated in Marco Colonna’s laboratory (Washington University).^43^ *Mda5*^−/−^ *Sting^Gt/Gt^* were bred in our lab.^44^ *CD64^cre^* (as known as *Fcgr1-cre)* mouse stain was generated in Dr. Ming O. Li’s laboratory (Memorial Sloan Kettering Cancer Center). *Ifnar1^fl/fl^* (Strain # 028256), *Mrp8^cre^* (Strain # 021614)^45,^, *LysM^cre^* (Strain #004781),^46^ and *CD4^cre^* (Strain #017336)^47^ mice were also purchased from Jackson Laboratory. *Mrp8^cre^Ifnar1^fl/fl^*, *Cd64^Cre^Ifnar1^fl/fl^*, *LysM^Cre^Ifnar1^fl/fl^*, *and Cd4^Cre^Ifnar1^fl/fl^* were bred in our lab.

### Cell lines

BHK-21 (baby hamster kidney cell, ATCC CCL-10) cells were cultured in Eagle’s minimal essential medium containing 10% fetal bovine serum (FBS), 0.1 mM nonessential amino acids, penicillin, and streptomycin. The murine melanoma cell line B16-F10 was originally obtained from I. Fidler (MD Anderson Cancer Center). MC38 cell line was obtained from ATCC. The B16-F10 cell line was maintained in RPMI-1640 medium supplemented with 10% FBS, 0.05 mM 2-mercaptoethanol, penicillin, and streptomycin. The B16-F10 cell line lacking B2M gene were generated by using CRISPR-cas9 technology. MB49-luc cell line was a kind gift from Michael Glickman (Memorial Sloan Kettering Cancer Center).

### Viruses

The MVA virus was a kind gift from Gerd Sutter (University of Munich). MVAΔE5R, MVAΔE5R-hFlt3LmOX40L (rMVA), rMVAΔE3L, rMVAΔE3LΔWR199 (MQ832), rMVAΔE3LΔWR199-IL12 (MQ833), and hMQ833 were generated by transfecting pUC57-based plasmids into BHK-21 cells that were infected with MVA at MOI 0.05. Recombinant viruses were purified after 4∼6 rounds of plaque selection based on the fluorescence marker. PCR and DNA sequencing were performed to verify the purity of the recombinant viruses. Viruses were propagated in BHK-21 cells and purified through a 36% sucrose cushion.

### Human monocyte-derived dendritic cell (moDC) generation and T cell isolation

All collection and use of human specimens adhered to protocols reviewed and approved by the Institutional Review and Privacy Board of Memorial Hospital, MSKCC. Buffy coat products were obtained from healthy donor at New York Blood Center and peripheral blood mononuclear cells (PBMCs) separated by standard centrifugation over Ficoll-Paque PLUS (Amersham Pharmacia Biotech, Uppsala, Sweden). Tissue culture plastic adherent PBMCs comprised the moDCs precursors, which were cultured in complete PRMI 1640-1% human serum supplemented with 1000 IU/ml GM-CSF (PeproTech) and 500 IU/ml IL-4 (PeproTech). Fresh medium and cytokines were replenished every 48 h. T cells were obtained from tissue culture nonadherent PBMCs, then further purified by CD8^+^ T cell or CD4^+^ T cell isolation kit (Miltenyi Biotec) following manufacturer’s instruction.

### Tumor processing and flow cytometry staining

Tumor samples were collected in cold RPMI and thoroughly chopped into small pieces with surgical scissor. Then we added Liberase TL (1.67 Wünsch U/ml) and DNase I (0.2 mg/ml) to the tube and incubated in 37 °C shaker for 20-30 minutes. Digested tumors were then transferred to GentleMACs C Tube and homogenized using the Gentle MACs Octo Dissociator (Miltenyi Biotec) according to manufacturer’s instruction. Samples were then filtered through 70 um filter and quenched with 10 ml cold PBS and spun down. Cells were then washed with MACS buffer (Miltenyi Biotec) twice, and stained with Fc block (BD), viability dye eFluor506 (eBioscience), and cell surface antibodies diluted in MACS buffer for 30 minutes on ice in the dark, and subsequently fixed and permeabilized using the Foxp3 fixation and permeabilization kit (Thermo Fisher). Cells were then incubated with intracellular antibodies diluted in permeabilization buffer for 30 minutes or overnight.

The following antibodies were used for flow cytometric staining: CD45 Pacific blue (clone 30-F11, Biolegend), CD3 BUV395 (clone 145-2C11, BD), CD4 BUV737 (clone RM4-5, BD), CD8 Alexa Fluor 700 (clone 53-6.7, Biolegend), OX40 BV605 (clone OX-86, Biolegend), TIM3 BV711 (clone RMT3-23, Biolegend), PD-1 AF647 (clone RMPI-30, BD), KLRG1 APC-Cy7(clone 2F1/KLRG1, Biolegend), Foxp3 PerCP-Cy5.5 (clone FJK-16s, Invitrogen), Ki67 BV786 (clone B56, BD), TOX PE (clone REA473, Miltenyi Biotec), Granzyme B PE-Texas red (clone GB11, Invitrogen), CD11b APC-eFluor780 (clone M1/70, eBioscience), Ly6G PE-Cy7 (clone 1A8, BD), F4/80 APC (clone BM8, Biolegend), CD68 BV421 (clone FA-11, Biolegend), iNOS PE (clone W16030C, Biolegend).

After staining, cells were washed and resuspended with MACS buffer and transferred into FACS tubes with 70 um filter caps. Single cell suspensions were run on Cytek Aurora analyzer, and the data was analyzed with Flowjo.

### Generation of bone marrow-derived dendritic cells

Bone marrow cells were extracted from the femur and tibia of mice. After centrifugation, cells were re-suspended in ACK Lysing Buffer (Lonza) for red blood cell lysis by incubating the cells on ice for 1-3 min. Cells were then resuspended in fresh medium and filtration through a 70-μm cell strainer (BD Biosciences). For the generation of Flt3L-BMDCs, the bone marrow cells (5 million cells in each well of 6-well plates) were cultured in complete medium (CM) in the presence of Flt3L (100 ng/ml; R & D systems) for 7–9 days. Cells were fed every 2–3 days by replacing 50% of the old medium with fresh medium. For the generation of GM-CSF-BMDCs, the bone marrow cells (5 million cells in each 15 cm cell culture dish) were cultured in RPMI-1640 medium supplemented with 10% fetal bovine serum (FBS) in the presence of 30 ng/ml GM-CSF (BioLegend) for 9-12 days. Fresh medium and cytokine were replenished every 2-3 days.

### RNA isolation and Real-time PCR

Cells were infected with various viruses at a MOI of 10 for 1 hour or mock-infected. The inoculum was removed, and the cells were washed with PBS twice and incubated with fresh medium for 16 hours or overnight. RNA was extracted from whole-cell lysates with RNeasy Plus Mini kit (Qiagen) and was reverse-transcribed with cDNA synthesis kit (Thermo Fisher). Real-time PCR was performed in triplicate with SYBR Green PCR Master Mix (Life Technologies) and Applied Biosystems 7500 Real-time PCR Instrument (Life Technologies) using gene-specific primers. Relative expression was normalized to the levels of glyceraldehyde-3-phosphate dehydrogenase (GAPDH). The primer sequences for quantitative real-time PCR are listed in <u>Table S1</u>.

### ELISA

For IL-12 transgene expression verification, B16 cells were infected with MVAΔE5R or MQ833 at a MOI of 10 for 1 hour. Mock-infected cells were used as a control. The supernatant was removed, and the cells were washed with PBS twice and incubated with fresh medium for 16 hours. Supernatant was collected from the culture, and IL-12 levels were determined using Duoset ELISA kit (R&D) following manufacturer’s instruction.

For T cell activation assays, supernatant from the co-culture system were collected and IFNγ levels were determined by Duoset ELISA kit (R&D).

### ELISPOT assay

Spleens were harvested from mice treated with different viruses and were mashed through a 70 µm strainer (Thermo Fisher Scientific). Red blood cells were lysed using ACK Lysis Buffer (Life Technology), and the cells were then resuspended in RPMI medium. Enzyme-linked ImmunoSpot (ELISPOT) assay was performed to measure IFN-γ+ CD8+ T cells, according to the manufacturer’s protocol (BD Bioscience). 1×10^6^ splenocytes were co-cultured with 2.5×10^5^ irradiated B16-F10 in complete RPMI medium overnight. IFNγ+ splenocytes were detected by Mouse IFNγ ELISPOT kit (BD Biosciences)

### Western blot analyses

B16-F10 cells or ^39^were infected with MQ832 or MQ833 at a MOI of 10 for 1 hour. Mock-infected cells were used as a control. Cells were then washed with PBS twice and cultured with fresh medium. Cells were lysed in RIPA lysis buffer supplemented with 1x Halt™ Protease and Phosphatase Inhibitor Cocktail at indicated time points. Protein samples were separated by SDS-PAGE and then transferred to nitrocellulose membrane. Primary antibodies specific for phospho-IRF3 (1:500, CST, 4947), IRF3 (1:1000, CST, 4302), cGAS (1:1000, CST, 31659), STING (1:1000, CST, 13647), phospho-STING (1:500, CST, 72971), TBK1 (1:1000, CST, 3504), and phosphor-TBK1 (1:500, CST, 5483) were used. GAPDH antibody (1:5000, CST, 2118) were used as loading controls. Anti-rabbit HRP-linked IgG antibody was used as a secondary antibody (1:5000, CST, 7074). Detection was performed using SuperSignalTM Substrates (Thermo Fisher, 34096 or A38555).

### In vitro T cell activation assay

For murine MQ833, splenic CD11c+ DCs were sorted from WT mice and subsequently infected with various viruses at a MOI of 10 for 1 hour and pulsed with OVA (1.25 mg/ml). OT-1 T cells were isolated from the spleens OT-1 mice using negative selection with CD8a^+^ T Cell Isolation Kit (Miltenyi Biotec) according to the manufacturer’s instructions. OT-1 T cells were then co-cultured with DCs (DC:T = 1:5) for 3 days. Alternatively, T cells were incubated with supernatant collected from the infected CD11c^+^ DCs for 3 days. IFN-γ levels in the supernatants were determined by ELISA.

For human MQ833, human moDCs were infected with various viruses at a MOI of 10 for 1 hour and pulsed with OVA (1.25 mg/ml). CD8 or CD4 T cells were purified from human PBMCs using isolation kits (Miltenyi Biotec) following manufacturer’s instruction. T cells and moDCs were co-cultured (DC:T = 1:15) for 3 days and supernatant were then collected for IFN-γ ELISA.

### Tumor challenge and treatment

For survival experiments, 5×10^5^ B16-F10 cells, 5×10^5^ MC38 cells, 1×10^6^ B2m^−/−^ B16 cells were implanted intradermally into the shaved skin on the right flanks of mice. 1×10^6^ MB49-luc cells were implanted subcutaneously into the right flanks of mice. Once the tumors are 3 mm in diameter or larger, they were injected with 4×10^7^ PFU of MQ832, MQ833 or PBS when the mice were under anesthesia. Viruses were injected twice weekly as specified in each experiment and tumor sizes were measured twice a week. For combination therapy with ICBs, the following antibodies were injected interperitoneally twice weekly: anti-CTLA-4 (100 µg per mouse), Anti-PD-L1 (200 µg per mouse), anti-PD-1 (200 per mouse), or isotype control antibody. Tumor volumes were calculated according to the following formula: l (length) × w (width) × h (height)/2. Mice were euthanized for signs of distress or when the diameter of the tumor reached 10 mm.

In the bilateral tumor implantation model, B16-F10 or MC38 cells were implanted intradermally into right (5×10^5^) and left (1×10^5^) flanks of C57BL/6J mice. After 7 days after implantation when the tumors were established, the tumors at the right flank were injected with 4×10^7^ PFU of MQ832, MQ833 or PBS twice weekly.

For the tumor rechallenge study, the survived mice (more than 60 days after initiation of intratumoral virotherapy) were rechallenged with intradermal delivery of a lethal dose of B16-F10, MB49 or B2m KO B16-F10 (5×10^5^ cells) at the contralateral side.

For flow cytometric analysis and single cell RNA-seq, 5×10^5^ B16-F10 tumor cells were implanted intradermally into the right flanks of the mice for the unilateral tumor implantation model. For bilateral implantation, 5×10^5^ and 2.5×10^5^ B16-F10 tumor cells were implanted intradermally into the right and left flanks of mice. At 8-11 days after implantation when tumors were established, the tumors at the right flank were injected with 4 ×10^7^ PFU of MQ832, MQ833, or PBS once, or twice with 3 days apart. Tumors were harvested two days after second injection.

### Purification of anti-CSF-1 monoclonal antibody

The anti-CSF-1 monoclonal antibody (clone 5A1) was purified from cell supernatant of hybridoma (ATCC, #CRL-2702) 1,2. Briefly, hybridoma cells were cultured in Hybridoma-SFM medium (Thermo #12045076) for 3 weeks to produce antibody. Cell supernatant was collected to do antibody purification by using LigaTrap Rat IgG purification column (LigaTrap Technologies #LT-138) according to the manufacturer’s procedure. Antibody quality and concentration were examined by SDS-PAGE gel electrophoresis and nanodrop, respectively.

### In vivo antibody depletion experiment

Immune cell subsets were depleted by administering 200 μg monoclonal antibodies i.p. twice weekly starting 1 day prior to the first viral injection as indicated, continued until the animals either died, were euthanized, or were completely clear of tumors: anti-CD8-α for CD8+ T cells (clone 2.43, BioXCell), anti-CD4 for CD4^+^ T cells (clone GK1.5, BioXCell), anti-NK1.1 for NK cells (clone PK136, BioXCell). 500 μg of anti-Ly6G (clone 1A8, BioXCell) were given for neutrophil depletion. 500 μg of anti-CSF1 (provided by Ming Li’s laboratory) were given every 5 days for macrophage depletion. Depletion efficacies of CD8 T cells, CD4 T cells, NK cells and neutrophils were confirmed by flow cytometry of peripheral blood.

### Cell sorting for tumor infiltrating immune cells

Tumors were harvested and single cell suspensions were prepared. Cells were then stained with Cell Viability Dye (eFlour506, brand) and anti-CD45 antibody (brand). CD45^+^ live cells were then sorted using B&D Aria cell sorter.

### Single-cell RNA-sequencing and data analysis

#### Library preparation and sequencing

CD45^+^ populations from tumors were purified by FACS sorting. We performed single-cell 5′ gene expression profiling on the single-cell suspension using the Chromium Single Cell V(D)J Solution from 10x Genomics according to the manufacturer’s instructions. Cell-barcoded 5′ gene expression libraries were sequenced on an Illumina NovaSeq6000 sequencer with pair-end reads.

#### Gene expression UMI counts matrix generation

The sequencing data were primarily analyzed by the 10× cellranger pipeline (v3.0.2) in two steps. In the first step, cellranger mkfastq demultiplexed samples and generated fastq files; and in the second step, cellranger count aligned fastq files to the reference genome and extracted gene expression UMI counts matrix. In order to measure both human and viral gene expression, we built a custom reference genome by integrating the MVA virus genome into the 10× pre-built mouse reference using cellranger mkref. The MVA virus genome was downloaded from NCBI.

#### Single-cell RNA-seq data analysis

All cells expressing <200 or >6,000 genes were removed as well as cells that contained >5% mitochondrial counts. Samples were merged and normalized. The default parameters of Seurat were used, unless mentioned otherwise. Briefly, 2,000 variable genes were identified for the clustering of all cell types and principal component analysis (PCA) was applied to the dataset to reduce dimensionality after regressing for the number of UMIs (counts). The top 12-15 most informative principal components (PCs) were used for clustering and Uniform Manifold Approximation and Projection for dimension reduction (UMAP). To characterize each cluster, we applied both the *FindAllMarkers* and *FindMarkers* procedure in Seurat, which identified markers using log fold changes (FC) of mean expression. To identify differentially expressed genes between two groups of clusters, *FindMarkers* functions in Seurat and Enhanced Volcano R package (v1.8.0) were used. For subsequent T cell analysis, we first extracted all the CD3^+^ cells from the original dataset and repeated the clustering procedures in the T cell subset.

#### Gene set enrichment analysis (GSEA)

The GSEA analysis was done and visualized using the *clusterProfiler* (v4.0.5) and *enrichplot* (v1.12.3) packages, which uses predefined gene sets from the Molecular Signatures Database (MSigDB v7.4) ^48^. We used the hallmark gene sets collection for the present study. Genes were ranked by the test statistic value obtained from differential expression analysis and the pre-ranked version of the tool was used to identify significantly enriched biological pathways.

## Statistical analysis

Two-tailed unpaired Student’s t test was used for comparisons of two groups in the studies. Survival data were analyzed by log-rank (Mantel-Cox) test. The p values deemed significant are indicated in the figures as follows: *, p < 0.05; **, p < 0.01; ***, p < 0.001; ****, p < 0.0001. The numbers of animals included in the study are discussed in each figure legend.

## Lead Contact

Further information and requests for resources and reagents used in this study should be directed to and will be fulfilled by the lead contact and the corresponding author, Liang Deng.

## Materials availability

Materials generated in our laboratory are available upon request.

## Data and code availability

The RNA-seq data reported in this study will be deposited in GEO and made publicly available prior to journal publication.

## Acknowledgements

We thank the Flow Cytometry Core Facility and Molecular Cytology Core Facility at the Sloan Kettering Institute and Genomic core at Weill Cornell Medical College. This work was supported by MSK Technology Development Fund (L.D.), Sponsored Research Agreement from IMVAQ Therapeutics (L.D.), Cycle for Survival (L.D.), Center for Experimental Therapeutics at Memorial Sloan Kettering Cancer Center (L.D.), Melanoma Research Alliance (L.D.), and SNF Institute for Global Infectious Disease Research (C.M.R.). This work was supported in part by the Swim across America (J.D.W., T.M.), Ludwig Institute for Cancer Research (J.D.W., T.M.), Parker Institute for Cancer Immunotherapy, Weill Cornell Medicine (J.D.W., T.M.). This research was also funded in part through the NIH/NCI Cancer Center Support Grant P30 CA008748.

## Author Contributions

S.T.L., Y.Q.W., and G.M. contributed to project conception, experimental design, execution of experiments, data analysis, interpretation, and manuscript writing. S.B.T., J.A.L., Y.W., and N.Y. assisted with mouse experiments and data analysis. S.T.L., Y.Q.W., T.Z., and A.T. analyzed the single-cell RNA-seq data. L.L.J. and M.O.L. provided the anti-CSF1 antibody and the CD64cre mouse strain. C.M.R., J.D.W., T.M., D.H., M.O.L., W.Y., J.C., J.H.W., and J.Z.X. provided input on experimental design, data interpretation, and manuscript editing. All authors contributed to manuscript preparation. L.D. supervised all aspects of the study.

