## Supplemental Figures for "Innate–adaptive IFN cross-talk reprograms myeloid and T cells for antitumor immunity with a second-generation recombinant MVA"

SUPPLEMENTAL ITEMS

This file contains:


- Figure. S1
- Figure. S2
- Figure. S3
- Figure. S4
- Figure. S5
- Figure. S6
- Figure. S7
- Figure. S8
- Figure. S9

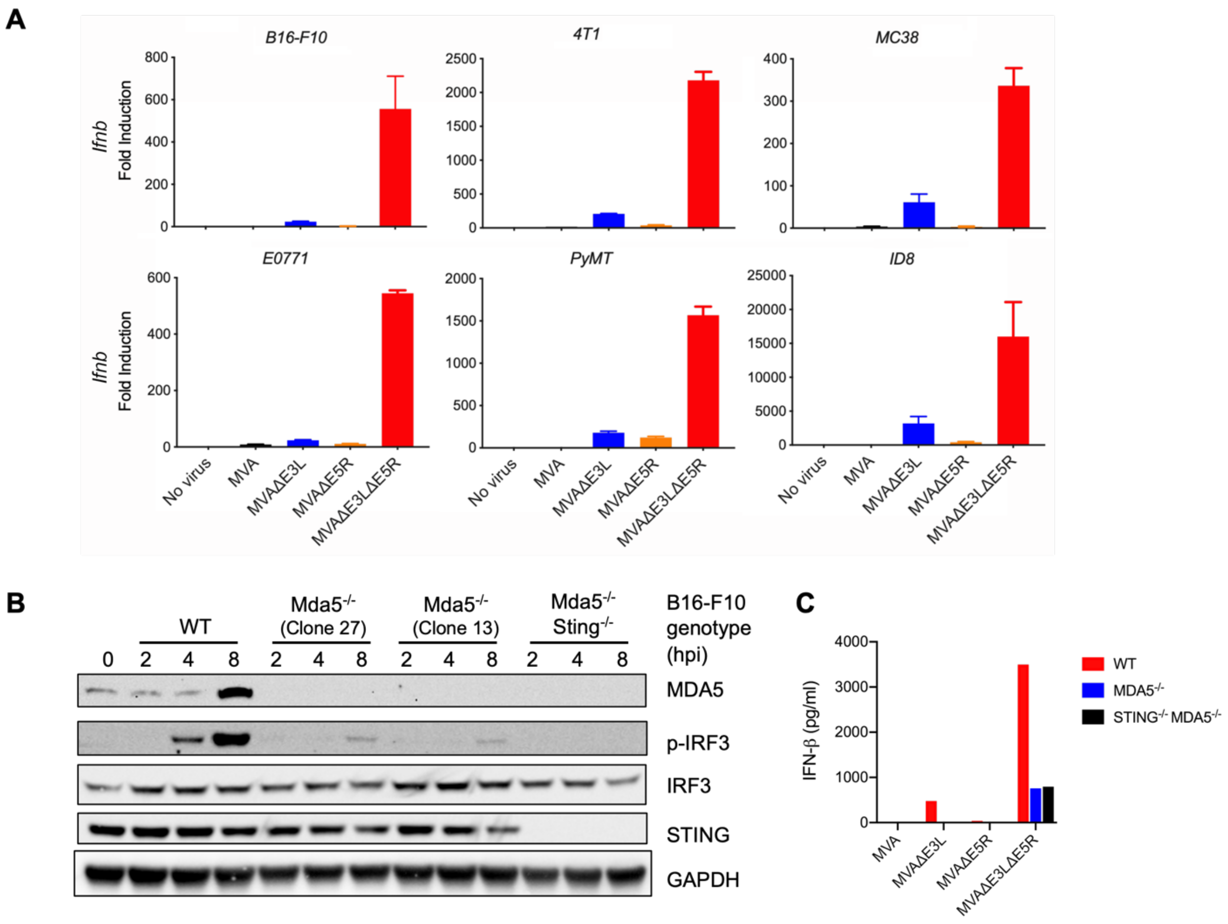
A
B16-F10
4T1
MC38
E0771
PyMT
ID8
Ifnb
Fold Induction
No virus
MVA
MVAΔE3L
MVAΔE5R
MVAΔE3LΔE5R
B
WT
Mda5-/- (Clone 27)
Mda5-/- (Clone 13)
Mda5-/- Sting-/-
B16-F10 genotype (hpi)
0 2 4 8 2 4 8 2 4 8 2 4 8
MDA5
p-IRF3
IRF3
STING
GAPDH
C
IFN-β (pg/ml)
WT
MDA5-/-
STING-/- MDA5-/-

**Figure S1. MVAΔE3LΔE5R promotes type I IFN expression in multiple tumor cell lines, related to Figure 1.**

(**A**) RT-PCR for *Ifnb* gene expression in B16-F10, 4T1, MC38, E0771, PyMT, and ID8 tumor cell lines with MVA, MVAΔE3L, MVAΔE5R, MVAΔE3LΔE5R virus infection.

(**B**) Western blot testing the expression of indicated proteins. WT, MDA5$^{-/-}$ (clone 27 and clone 13), and MDA5$^{-/-}$STING$^{-/-}$ B16-F10 cells were infected with MVAΔE3LΔE5R viruses at a MOI of 10, and cells were collected at 2, 4, 8 hours post infection (hpi).

(**C**) ELISA test for IFN-β production. WT, MDA5$^{-/-}$, and MDA5$^{-/-}$STING$^{-/-}$ B16-F10 cells were infected by MVAΔE5R or MQ833 at a MOI of 10. Supernatant were collected at 16 hours post infection (hpi).

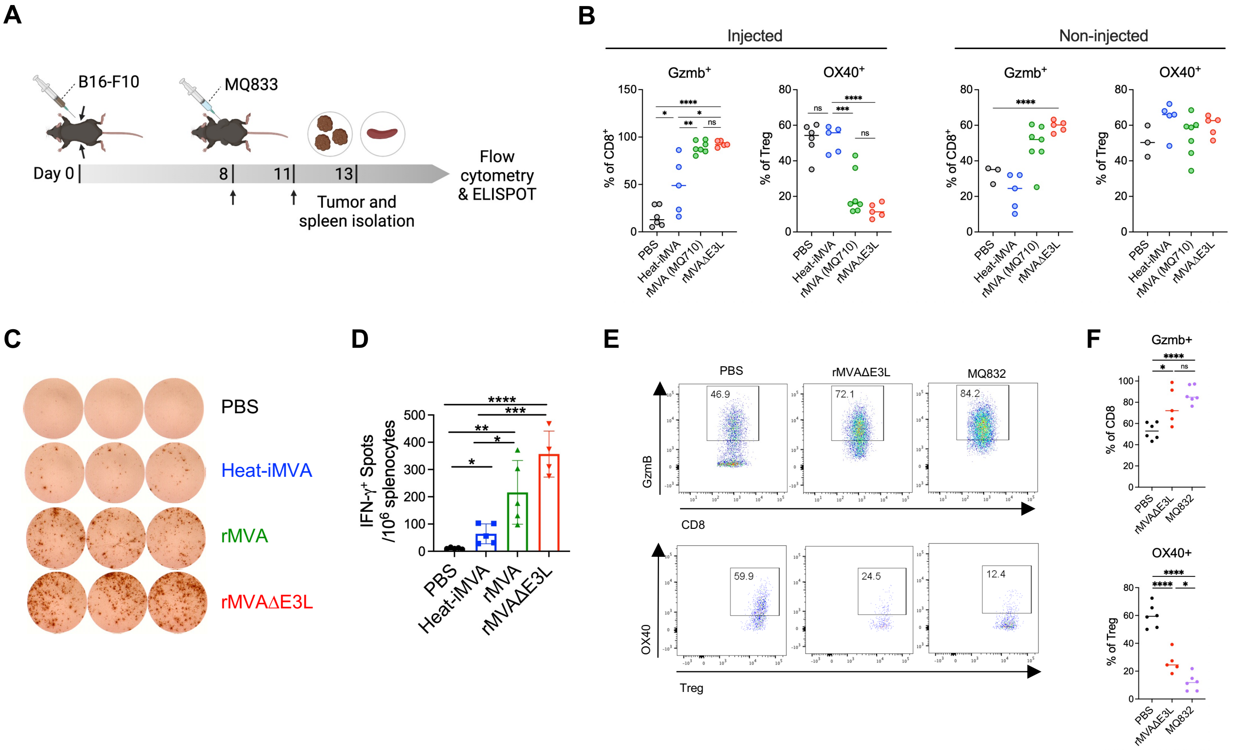
A
B16-F10
MQ833
Day 0
8
11
13
Tumor and spleen isolation
Flow cytometry & ELISPOT
B
Injected
Non-injected
Gzmb+
OX40+
% of CD8+
% of Treg
PBS
Heat-iMVA
rMVA (MQ710)
rMVAΔE3L
C
PBS
Heat-iMVA
rMVA
rMVAΔE3L
D
IFN-γ+ Spots /10^6 splenocytes
E
PBS
rMVAΔE3L
MQ832
46.9
72.1
84.2
GzmB
CD8
59.9
24.5
12.4
OX40
Treg
F
Gzmb+
% of CD8
OX40+
% of Treg
MQ832

**Figure S2. Antitumor activities generated by IT rMVAΔE3L and rMVAΔE3LΔWR199 (MQ832), related to Figure 1.**

(**A**) Diagram of the experimental design.

(**B**) Dot plots showing the percentages of Gzmb$^+$ cells among CD8$^+$ T cells (left) and OX40$^+$ cells among CD45$^+$CD3$^+$CD4$^+$Foxp3$^+$ Tregs in B16-F10 tumors treated with Heat-iMVA, rMVA, or rMVAΔE3L using a flow cytometric analysis.

(**C-D**) ELISPOT assay analyzing CD8$^+$ T cells isolated from splenocytes of mice treated with different viruses for anti-tumor IFN-$\gamma^+$ T cells. (**C**) Representative images from the ELISPOT assay. (**D**) IFN-$\gamma^+$ spots per 1,000,000 purified CD8$^+$ T cells from the spleens of the mice treated with IT PBS, Heat-iMVA, rMVA, or rMVAΔE3L (n=5, $*P < 0.05$; $**P < 0.01$; $***P < 0.001$; $****P < 0.0001$).

(**E-F**) Flow cytometric analysis of tumor infiltrating T cells in virus treated B16-F10 tumors. (**E**) Dot plots showing percentages of Gzmb$^+$ cells among CD8 T cells (top) and OX40$^+$ cells among CD45$^+$CD3$^+$CD4$^+$Foxp3$^+$ Tregs in B16-F10 tumors treated with PBS, rMVAΔE3L, or rMVAΔE3LΔWR199 (MQ833). (**F**) Percentages of CD8$^+$ T cells expressing GzmB (top) and Tregs expressing OX40 (bottom) within tumors from different treatment groups. (n=5-6, $*P < 0.05$; $**P < 0.01$; $***P < 0.001$; $****P < 0.0001$).

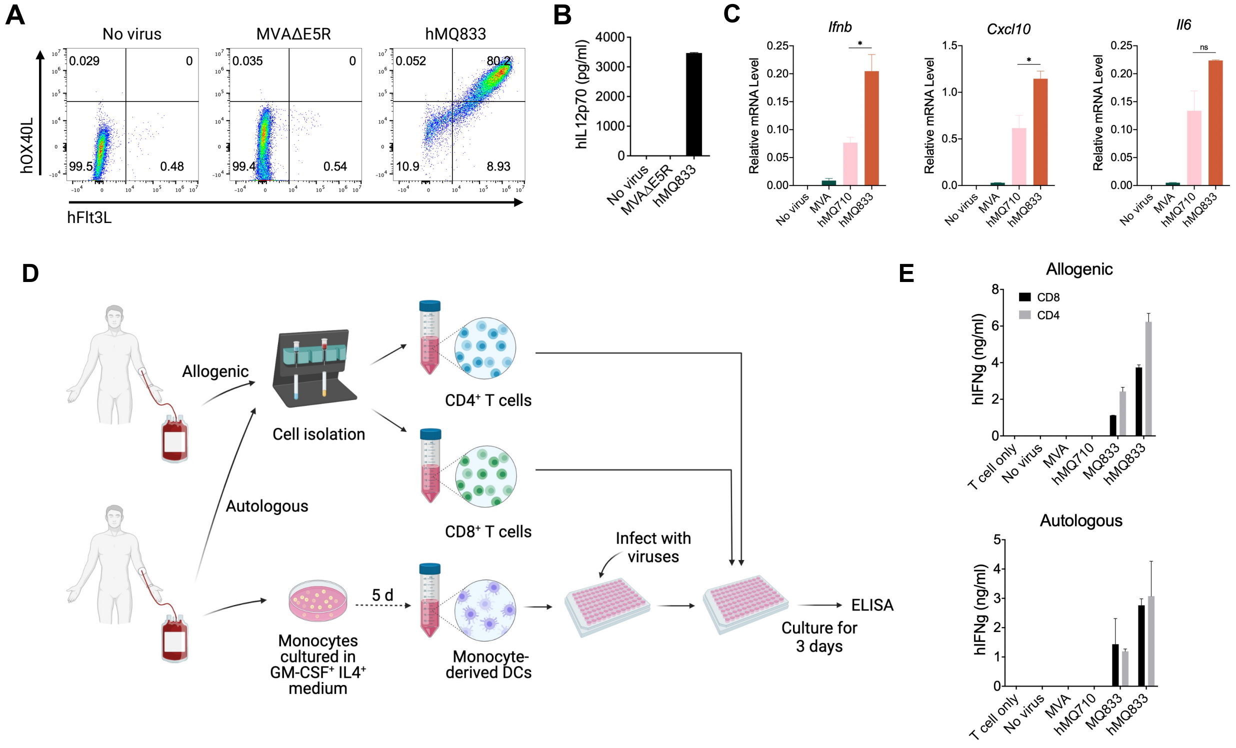

A
No virus
MVAΔE5R
hMQ833
hOX40L
hFlt3L
0.029
0
99.5
0.48
0.035
0
99.4
0.54
0.052
80.2
10.9
8.93
B
hIL12p70 (pg/ml)
No virus
MVAΔE5R
hMQ833
C
Ifnb
Cxcl10
Il6
Relative mRNA Level
No virus
MVA
hMQ710
hMQ833
ns
D
Allogenic
Autologous
Cell isolation
CD4⁺ T cells
CD8⁺ T cells
Monocytes cultured in GM-CSF⁺ IL4⁺ medium
5 d
Monocyte-derived DCs
Infect with viruses
Culture for 3 days
ELISA
E
Allogenic
Autologous
CD8
CD4
hIFNg (ng/ml)
T cell only
No virus
MVA
hMQ710
MQ833
hMQ833

**Figure S3: MQ833 promotes T cell activation by DCs in human PBMCs. Related to Figure 1.**

(**A-B**) Human monocyte-derived dendritic cells (MoDCs) were infected with indicated viruses at a MOI of 10. Cells and supernatant were collected 16 hours post infection. (A) Flow cytometric analysis of hOX40L and hFlt3L transgene expression. (B) ELISA assay for hIL12p70.

(**C**) qRT-PCR analyses of *Ifnb*, *Cxcl10*, and *Il6* gene expression on MoDCs infected with various viruses.

(**D**) Schematic diagram for human MoDC and T cell coculture experiment. MoDCs were isolated and cultured from the human PBMCs, pulsed with OVA, and infected with different viruses. Allogenic or autologous CD8 or $CD4^+$ T cells were isolated from PBMC of a different or the same donor, respectively, and co-cultured with moDCs. Supernatants were collected and IFN-γ level was measured by ELISA.

(**E**) ELISA for IFN-γ level in supernatants from the coculture system.

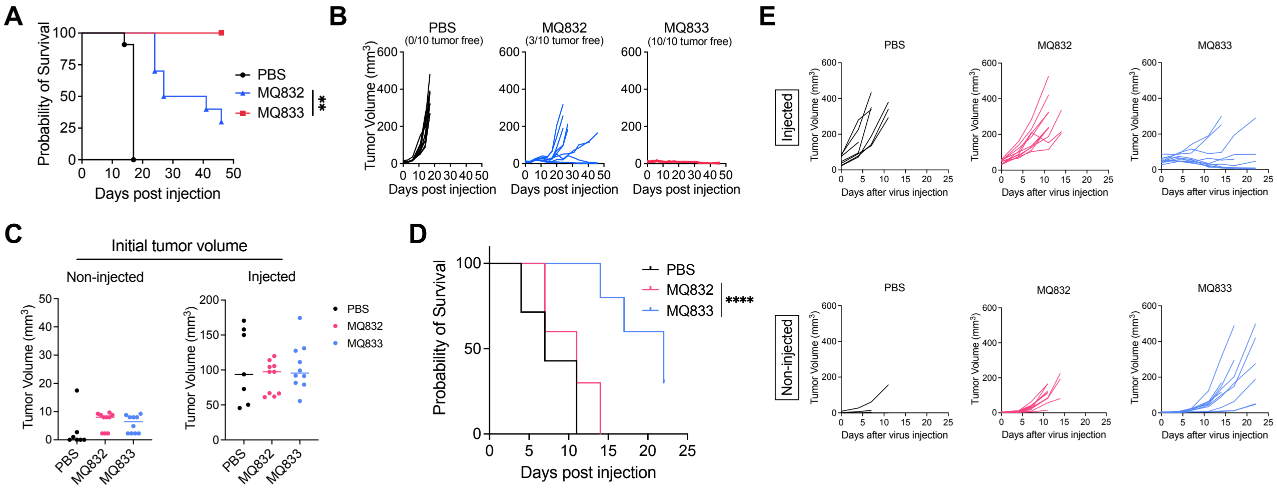
A
Probability of Survival
Days post injection
PBS
MQ832
MQ833
**
B
PBS (0/10 tumor free)
MQ832 (3/10 tumor free)
MQ833 (10/10 tumor free)
Tumor Volume (mm3)
Days post injection
C
Initial tumor volume
Non-injected
Injected
Tumor Volume (mm3)
PBS
MQ832
MQ833
D
Probability of Survival
Days post injection
PBS
MQ832
MQ833
****
E
Injected
Non-injected
PBS
MQ832
MQ833
Tumor Volume (mm3)
Days after virus injection

**Figure S4: IT MQ833 generates strong antitumor response in MC38 model. Related to Figure 2.**

(A-B) 5x10$^{5}$ MC38 cells were implanted intradermally into the right flank of WT C57BL/6J mice. Intratumoral injections of MQ832, MQ833, or PBS were given twice weekly once the tumors were established. Tumor volumes and mice survival were monitored. (A) Kaplan-Meier survival curve of mice in each treatment group. PBS was used as a control. (n=10) Survival data were analyzed by log-rank (Mantel-Cox) test. (*, $P < 0.05$). (B) Tumor volumes of mice separated by group over days post treatment.

(C-E) MC38 cells were implanted intradermally into the left (1x10$^{5}$ cells) and right (5x10$^{5}$ cells) flanks of WT C57BL/6J mice. Intratumoral injections of MQ832, MQ833, or PBS were given twice weekly once the injected tumors reaches 5 mm in diameter. (C) Initial tumor volumes upon first virus injection in each group. (D) Kaplan-Meier survival curve of mice in each treatment group. PBS was used as a control. (n=10) Survival data were analyzed by log-rank (Mantel-Cox) test (****$P < 0.0001$). (E) Injected (top) and non-injected (bottom) tumor volumes of mice separated by treatment groups over days post treatment.

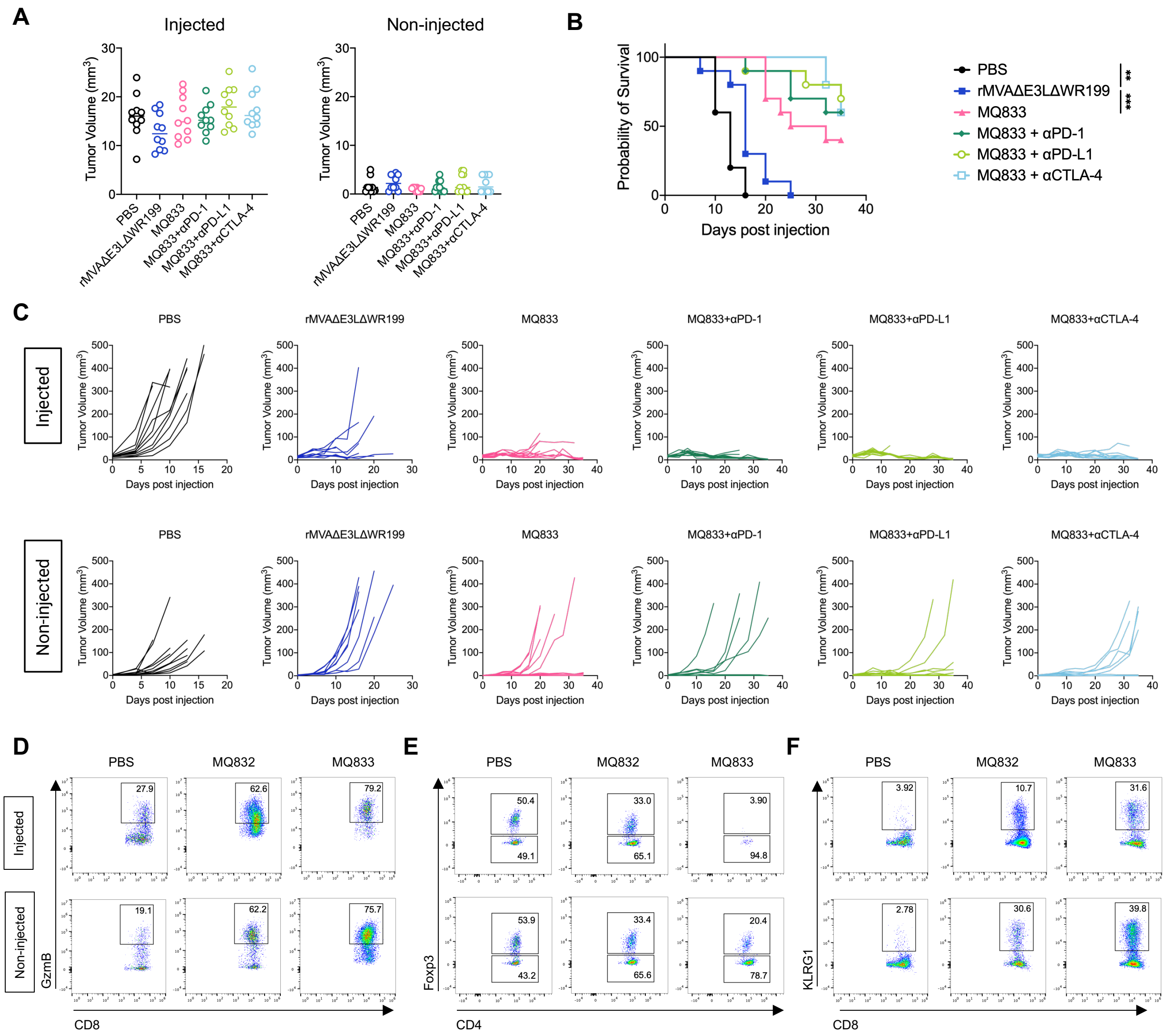
A
Injected
Non-injected
Tumor Volume (mm³)
PBS
rMVAΔE3LΔWR199
MQ833
MQ833+αPD-1
MQ833+αPD-L1
MQ833+αCTLA-4
B
Probability of Survival
Days post injection
PBS
rMVAΔE3LΔWR199
MQ833
MQ833 + αPD-1
MQ833 + αPD-L1
MQ833 + αCTLA-4
**
***
C
Injected
Non-injected
Days post injection
D
E
F
PBS
MQ832
MQ833
GzmB
CD8
Foxp3
CD4
KLRG1
27.9
62.6
79.2
19.1
62.2
75.7
50.4
49.1
33.0
65.1
3.90
94.8
53.9
43.2
33.4
65.6
20.4
78.7
3.92
10.7
31.6
2.78
30.6
39.8

**Figure S5: IT MQ833 generates abscopal antitumor effects facilitated by combination therapy with ICBs. Related to Figure 2.**

(**A-C**) B16-F10 cells were implanted intradermally into the left ($1x10^5$ cells) and right ($5x10^5$ cells) flanks of WT C57BL/6J mice. Intratumoral injections of rMVAΔE3LΔWR199 (MQ832), MQ833, or PBS were given twice weekly once tumors were established on both sides. αPD-1, αPD-L1, or αCTLA-4 antibodies were given i.p. in combination of the MQ833 injection. Tumor volumes and mice survival were monitored.

(**D-F**) WT C57BL/6J mice were implanted intradermally with B16-F10 cells on the right ($5x10^5$) and left ($2.5x10^5$) flanks. Once the tumors were established, 2 doses of $4x10^7$ PFU of MQ833 were intratumorally delivered to the right flanks twice, 3 days apart. Injected and non-injected tumors were harvested at day 2 post the second injection and then stained and analyzed by flow cytometry. (**D**) Representative dot plots showing percentages of $CD8^+$ T cells expressing Gzmb, (**E**) $CD4^+$ T cells expressing Foxp3 (Tregs) and (**F**) $CD8^+$ T cells expressing KLRG1 in the injected (top) and non-injected (bottom) tumors.

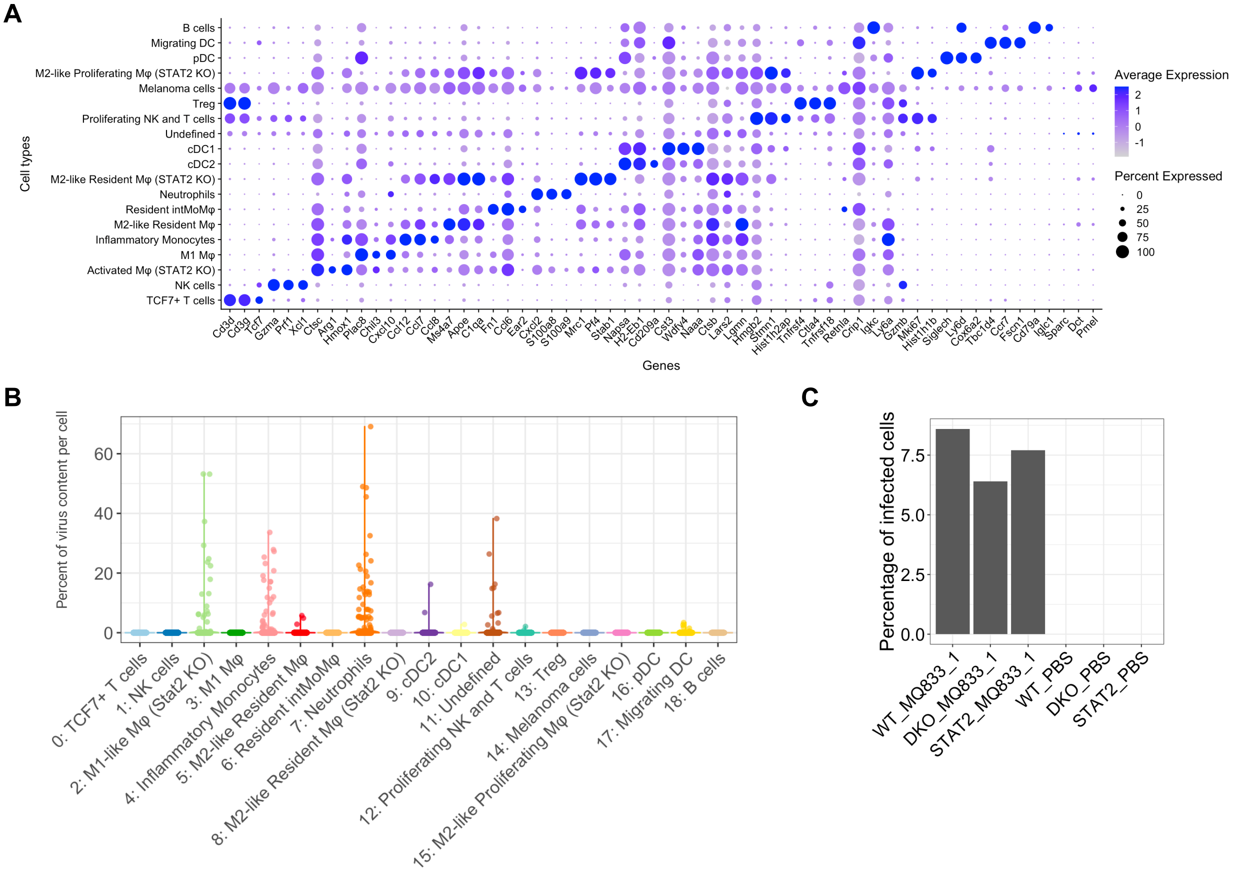
A
B cells
Migrating DC
pDC
M2-like Proliferating Mφ (STAT2 KO)
Melanoma cells
Treg
Proliferating NK and T cells
Undefined
cDC1
cDC2
M2-like Resident Mφ (STAT2 KO)
Neutrophils
Resident intMoMφ
M2-like Resident Mφ
Inflammatory Monocytes
M1 Mφ
Activated Mφ (STAT2 KO)
NK cells
TCF7+ T cells
Cell types
Cd3d Cd3g Tcf7 Gzma Prf1 Xcl1 Ctsc Arg1 Hmox1 Plac8 Chil3 Cxcl10 Ccl12 Ccl7 Ccl8 Ms4a7 Apoe C1qa Fn1 Ccl6 Ear2 Cxcl2 S100a8 S100a9 Mrc1 Pf4 Stab1 Napsa H2-Eb1 Cd209a Cst3 Wdfy4 Naaa Ctsb Lars2 Lgmn Hmgb2 Stmn1 Hist1h2ap Tnfrsf4 Ctla4 Tnfrsf18 Retnla Crip1 Igkc Ly6a Gzmb Mki67 Hist1h1b Siglech Ly6d Cox6a2 Tbc1d4 Ccr7 Fscn1 Cd79a Iglc1 Sparc Dct Pmel
Genes
Average Expression
2
1
0
-1
Percent Expressed
0
25
50
75
100
B
Percent of virus content per cell
0
20
40
60
0: TCF7+ T cells
1: NK cells
2: M1-like Mφ (Stat2 KO)
3: M1 Mφ
4: Inflammatory Monocytes
5: M2-like Resident Mφ
6: Resident intMoMφ
7: Neutrophils
8: M2-like Resident Mφ (Stat2 KO)
9: cDC2
10: cDC1
11: Undefined
12: Proliferating NK and T cells
13: Treg
14: Melanoma cells
15: M2-like Proliferating Mφ (Stat2 KO)
16: pDC
17: Migrating DC
18: B cells
C
Percentage of infected cells
0.0
2.5
5.0
7.5
WT_MQ833_1
DKO_MQ833_1
STAT2_MQ833_1
WT_PBS
DKO_PBS
STAT2_PBS

**Figure S6: scRNA-seq analysis of tumor infiltrating immune cells after IT MQ833 injection. Related to Figure 3.**

(**A-C**), $5x10^5$ of B16-F10 cells were intradermally implanted into the right flank of WT C57BL/6J mice, *Mda5*$^{-/-}$ *Sting*$^{Gt/Gt}$, or *Stat2*$^{-/-}$ mice. One dose of IT MQ833 injection was given once the tumors were established. Tumors were collected and processed into single cell suspension, and $CD45^+$ cells were sorted. scRNA-seq was performed. (**A**) Dot plots of representative differentially expressed genes for individual clusters. (**B**) Dot plot showing percentages of viral gene transcripts per cell in each cluster. One dot represents one cell. (**C**) Bar plot showing percentage of virus infected cells (virus transcript > 0) of total tumor infiltrating $CD45^+$ cells in each treatment group.

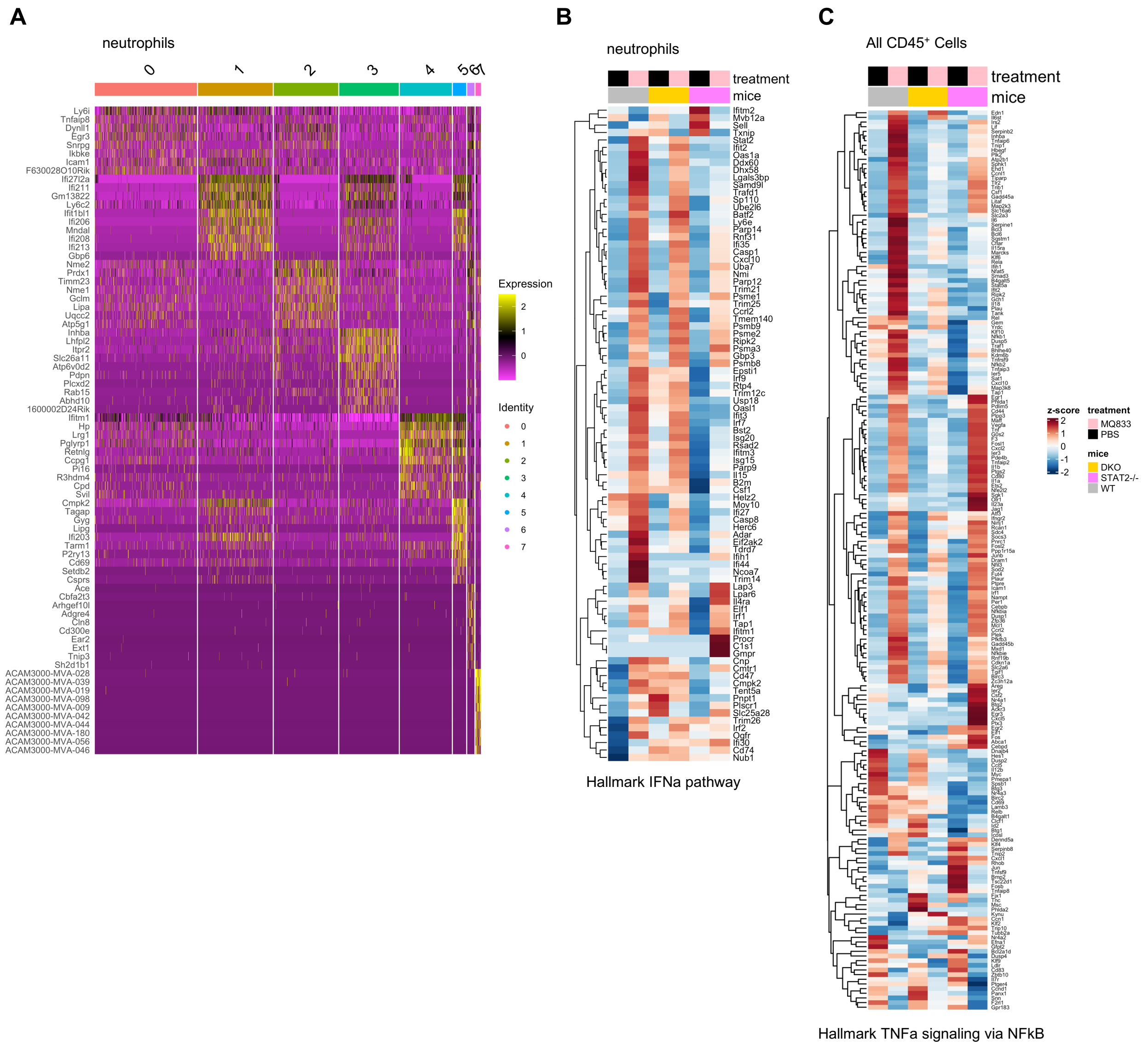
A
neutrophils
0
1
2
3
4
5
6
7
Ly6i
Tnfaip8
Dynll1
Egr3
Snrpg
Ikbke
Icam1
F630028O10Rik
Ifi27l2a
Ifi211
Gm13822
Ly6c2
Ifit1bl1
Ifi206
Mndal
Ifi208
Ifi213
Gbp6
Nme2
Prdx1
Timm23
Nme1
Gclm
Lipa
Uqcc2
Atp5g1
Inhba
Lhfpl2
Itpr2
Slc26a11
Atp6v0d2
Pdpn
Plcxd2
Rab15
Abhd10
1600002D24Rik
Ifitm1
Hp
Lrg1
Pglyrp1
Retnlg
Ccpg1
Pi16
R3hdm4
Cpd
Svil
Cmpk2
Tagap
Gyg
Lipg
Ifi203
Tarm1
P2ry13
Cd69
Setdb2
Csprs
Ace
Cbfa2t3
Arhgef10l
Adgre4
Cln8
Cd300e
Ear2
Ext1
Tnip3
Sh2d1b1
ACAM3000-MVA-028
ACAM3000-MVA-039
ACAM3000-MVA-019
ACAM3000-MVA-098
ACAM3000-MVA-009
ACAM3000-MVA-042
ACAM3000-MVA-044
ACAM3000-MVA-180
ACAM3000-MVA-056
ACAM3000-MVA-046
Expression
2
1
0
Identity
0
1
2
3
4
5
6
7
B
neutrophils
treatment
mice
Ifitm2
Mvb12a
Sell
Txnip
Stat2
Ifit2
Oas1a
Ddx60
Dhx58
Lgals3bp
Samd9l
Trafd1
Sp110
Ube2l6
Batf2
Ly6e
Parp14
Rnf31
Ifi35
Casp1
Cxcl10
Uba7
Nmi
Parp12
Trim21
Psme1
Trim25
Ccrl2
Tmem140
Psmb9
Psme2
Ripk2
Psma3
Gbp3
Psmb8
Epsti1
Irf9
Rtp4
Trim12c
Usp18
Oasl1
Ifit3
Irf7
Bst2
Isg20
Rsad2
Ifitm3
Isg15
Parp9
Il15
B2m
Csf1
Helz2
Mov10
Ifi27
Casp8
Herc6
Adar
Eif2ak2
Tdrd7
Ifih1
Ifi44
Ncoa7
Trim14
Lap3
Lpar6
Il4ra
Elf1
Irf1
Tap1
Ifitm1
Procr
C1s1
Gmpr
Cnp
Cmtr1
Cd47
Cmpk2
Tent5a
Pnpt1
Plscr1
Slc25a28
Trim26
Irf2
Ogfr
Ifi30
Cd74
Nub1
Hallmark IFNa pathway
C
All CD45+ Cells
treatment
mice
z-score
2
1
0
-1
-2
treatment
MQ833
PBS
mice
DKO
STAT2-/-
WT
Hallmark TNFa signaling via NFkB

**Figure S7: scRNA-seq analysis of neutrophil subclusters and CD45$^+$ myeloid cells. Related to Figure 4.**

(**A**) Heatmap of the average expression of the top 10 differentially expressed genes in each neutrophil subcluster

(**B**) Heatmap of the average expression of genes in the hallmark IFNα response gene set in neutrophil subcluster

(with gene names, related to Figure 4G)

(**C**) Heatmap of the average expression of genes in the Hallmark TNFa signaling via NFkB in all CD45 cells

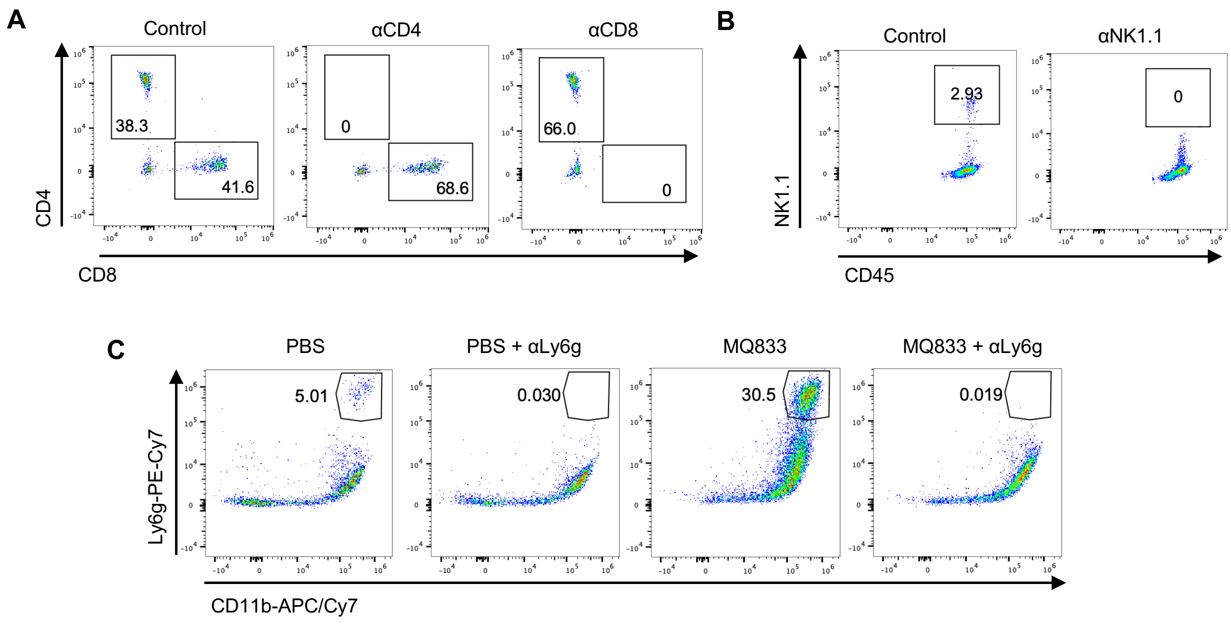
A
Control
αCD4
αCD8
38.3
41.6
0
68.6
66.0
0
CD4
CD8
B
Control
αNK1.1
2.93
0
NK1.1
CD45
C
PBS
PBS + αLy6g
MQ833
MQ833 + αLy6g
5.01
0.030
30.5
0.019
Ly6g-PE-Cy7
CD11b-APC/Cy7

**Figure S8: Flow cytometric verification of antibody depletion efficacy. Related to Figure 6.**

Peripheral blood were collected from mice treated with different depletion antibodies. Depletion of specific immune cell population was measured by fluorescent staining and flow cytometry.

(**A**) Representative dot plots showing percentages of CD8$^+$ and CD4$^+$ T cells from tumors injected with no antibody (left), αCD4 antibody (middle), and αCD8 antibody.

(**B**) Representative dot plots showing percentages of NK1.1$^+$ NK cells from tumors injected with no antibody (left) and αNK1.1 antibody.

(**C**) Representative dot plots showing percentages of Ly6g$^+$CD11b$^+$ neutrophils from tumors injected with PBS or MQ833 in combination of αLy6g antibody.

(**D**) Tumor volumes of indicated groups over days post injection

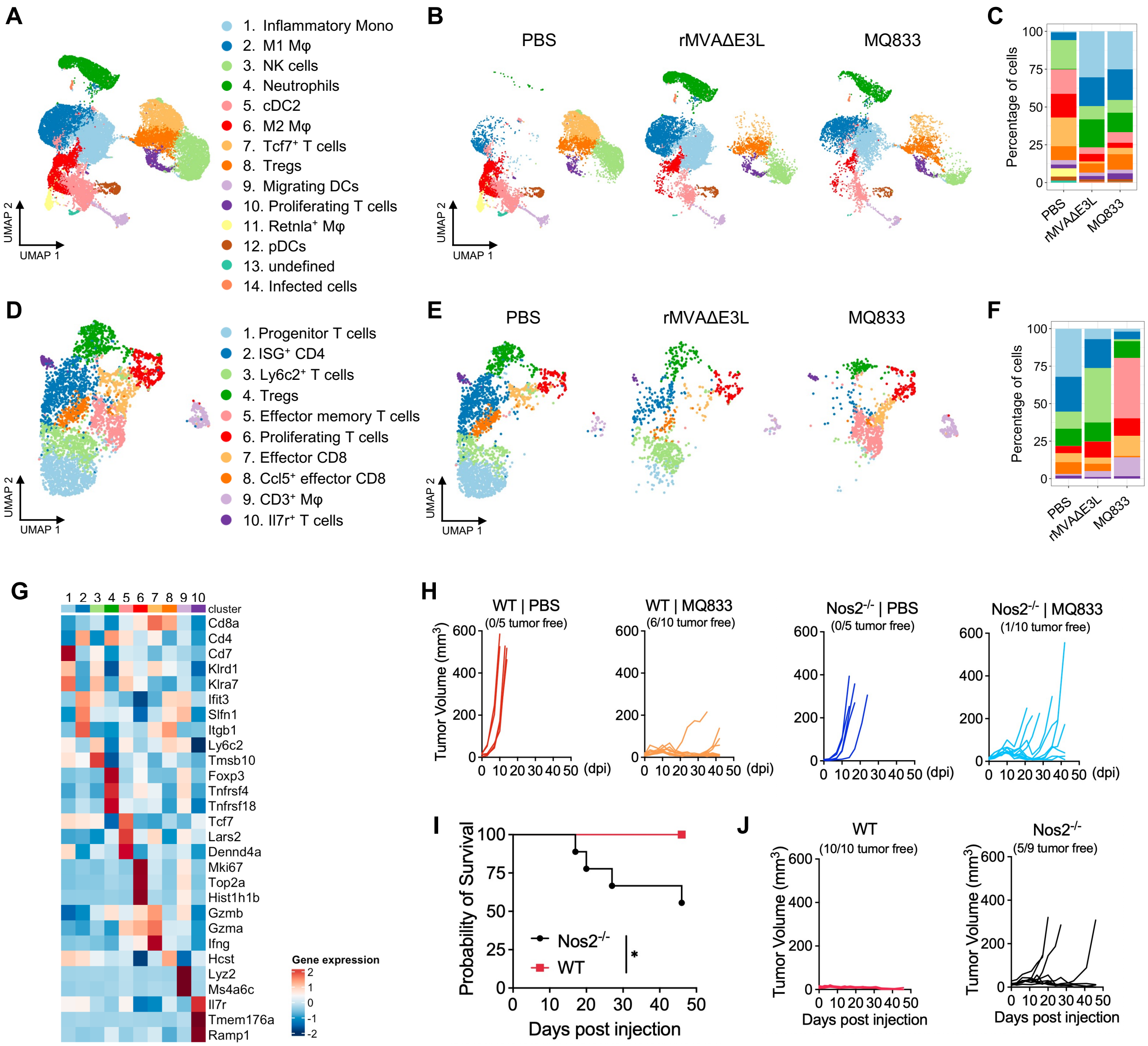
A
1. Inflammatory Mono
2. M1 Mφ
3. NK cells
4. Neutrophils
5. cDC2
6. M2 Mφ
7. Tcf7+ T cells
8. Tregs
9. Migrating DCs
10. Proliferating T cells
11. Retnla+ Mφ
12. pDCs
13. undefined
14. Infected cells
UMAP 1
UMAP 2
B
PBS
rMVAΔE3L
MQ833
C
Percentage of cells
D
1. Progenitor T cells
2. ISG+ CD4
3. Ly6c2+ T cells
4. Tregs
5. Effector memory T cells
6. Proliferating T cells
7. Effector CD8
8. Ccl5+ effector CD8
9. CD3+ Mφ
10. Il7r+ T cells
E
F
G
cluster
Cd8a
Cd4
Cd7
Klrd1
Klra7
Ifit3
Slfn1
Itgb1
Ly6c2
Tmsb10
Foxp3
Tnfrsf4
Tnfrsf18
Tcf7
Lars2
Dennd4a
Mki67
Top2a
Hist1h1b
Gzmb
Gzma
Ifng
Hcst
Lyz2
Ms4a6c
Il7r
Tmem176a
Ramp1
Gene expression
H
WT | PBS
(0/5 tumor free)
WT | MQ833
(6/10 tumor free)
Nos2-/- | PBS
(0/5 tumor free)
Nos2-/- | MQ833
(1/10 tumor free)
Tumor Volume (mm3)
(dpi)
I
Probability of Survival
Nos2-/-
WT
*
Days post injection
J
WT
(10/10 tumor free)
Nos2-/-
(5/9 tumor free)
Tumor Volume (mm3)
Days post injection

**Figure S9: Single cell RNA-seq analysis of tumor infiltrating CD45$^+$ cells after rMVAΔE3L and MQ833 treatment. Related to Figure 7.**

(**A-G**) 5x10$^5$ B16-F10 cells were intradermally implanted into the right flank of WT C57BL/6J mice. Once the tumors were established, rMVAΔE3L, MQ833, or PBS injection were delivered intratumorally. Tumors were collected 2 days after the injection. CD45$^+$ cells were sorted and sent for scRNA-seq. (**A**) UMAP display of sorted CD45$^+$ cells from 3 samples combined following 10X Genomics scRNA-seq workflow (n = 28,223 cells). (**B**) UMAP plots of cell clusters separated by sample. (**C**) Stacked bar plot showing the percentage of each cluster across different samples. (**D**) UMAP display of subclustered CD45$^+$CD3$^+$ cells from 6 samples combined following 10X Genomics scRNA-seq workflow (n = 4,618 cells). (**E**) UMAP plots of cell clusters separated by sample. (**F**) Stacked bar plot showing the percentage of each cluster across different samples. (**G**) Heatmaps of the average expression of representative marker genes for each cluster. (**H**) Tumor volumes of WT or *Nos2*$^{-/-}$ mice separated by group over days post treatment. 5x10$^5$ of B16-F10 cells were intradermally implanted into the right flank of *Nos2*$^{-/-}$ mice. IT MQ833 or PBS injections were given twice a week once the tumors were established. (**I-J**) 5x10$^5$ MC38 cells were implanted intradermally into the right flank of WT C57BL/6J and *Nos2*$^{-/-}$ mice. Intratumoral injections of MQ833 were given twice weekly once the tumors were established. Tumor volumes and mice survival were monitored. (**I**) Kaplan-Meier survival curve of mice in each treatment group (WT: n=10, *Nos2*$^{-/-}$, n = 9). Survival data were analyzed by log-rank (Mantel-Cox) test. (**, P < 0.05*). (**J**) Tumor volumes of mice separated by group over days post treatment.
